# An integrated landscape of mRNA and protein isoforms

**DOI:** 10.1101/2025.04.16.648713

**Authors:** Amir Kedan, Henrik Zauber, Meng-Ran Wang, Qionghua Zhu, Liang Fang, Wei Chen, Matthias Selbach

## Abstract

Alternative splicing and proteolytic processing expand the functional diversity of the human proteome by generating distinct protein isoforms from a single gene. However, the extent to which transcript isoforms give rise to distinct protein products remains unclear, in part due to technical limitations in proteomic workflows. Here, we combine full-length mRNA sequencing with SDS-PAGE-based protein fractionation and quantitative mass spectrometry to construct an integrated landscape of mRNA and protein isoforms in human RPE-1 cells. To overcome the inherent ambiguity of bottom-up proteomics in isoform detection, we developed *IsoFrac*, a computational pipeline that resolves protein isoforms based on their migration profiles across gel fractions. This approach enabled the identification of ∼45,000 full-length transcripts, ∼32,000 ORFs, and ∼16,000 distinct protein isoforms. Comparative analyses revealed widespread translation of alternative transcripts and uncovered proteolytic processing as a major, underappreciated source of proteome complexity. Our results establish a scalable framework for isoform-resolved proteogenomics and provide a reference resource for studying the molecular diversity encoded by the human genome.

## Introduction

Gene expression is a complex and tightly regulated process that transforms the genetic information into functional proteins ^1,2^. This process begins with the transcription of genes into mRNAs and their subsequent translation into proteins and is further diversified by the generation of multiple mRNA and protein isoforms from individual genes. Posttranscriptional processes, such as alternative splicing and alternative polyadenylation, give rise to diverse mRNA species ^3–5^. On the protein level, additional layers of complexity emerge through mechanisms like alternative translation initiation and posttranslational modifications ^6–8^. It is estimated that an average cell line may contain over one million distinct proteoforms, though this number varies by cell type and tissue.

While long-read sequencing has provided a detailed view of transcriptome diversity, the global characterization of protein isoforms remains incomplete. A persistent challenge in the field is the consistent discrepancy between the number of mRNA isoforms detected by RNA sequencing and the number of protein isoforms identified by proteomics. This raises two key questions: (1) To what extent do alternative transcripts yield functional protein products, and (2) how many additional protein isoforms arise through mechanisms independent of mRNA variation, such as alternative translation initiation or proteolytic processing? Two competing hypotheses have been proposed to explain the first question: One suggests that many alternative transcripts are not translated and may represent non-functional or regulatory RNA species ^9,10^, while the other posits that most splice variants are translated and actively contribute to proteome diversity ^3,5^. However, an alternative explanation is that technical limitations in proteomic methodologies prevent the reliable detection of many protein isoforms. To resolve this issue, it is essential to bridge the gap between transcript-level predictions and protein isoform identification, addressing both biological and technical constraints ^6^. Here, we refer to *’protein isoforms*’ as distinct polypeptides arising from different transcript isoforms, alternative translation or proteolytic cleavage events. This contrasts with ’*proteoforms*’, a broader term that includes all molecular forms of a protein, including those generated by post-translational modifications ^7^.

Current mass spectrometry (MS)-based approaches are typically divided into Top-Down and Bottom-Up methods, each facing distinct technical challenges in protein isoform identification. The Top-Down approach directly analyzes intact proteins, making it well-suited for isoform identification ^11^. However, the extreme physicochemical diversity of proteins complicates comprehensive proteome coverage, and despite recent advances, Top-Down proteomics still lags behind the depth achieved by Bottom-Up techniques ^12,13^. In contrast, the Bottom-Up approach analyzes peptides derived from digested proteins, achieving greater proteomic depth—often exceeding 10,000 protein groups in a single experiment ^14^. However, its ability to distinguish individual protein isoforms is limited, as many peptides are shared among multiple isoforms, making precise assignments challenging ^15^.

Most efforts to overcome the limitations of protein isoform identification have focused on deeper proteome sequencing and advanced data processing algorithms to extract isoform information from existing datasets ^16–20^. For example, a recent study employing six different proteases and three tandem mass spectrometry fragmentation methods in a deep Bottom-Up proteomics approach identified approximately one-third of observed alternative splice variants at the protein level and up to 64% among high abundant transcripts ^21^. Recent work has also demonstrated that thermal proteome profiling combined with deep peptide coverage can be used to cluster peptides into functional proteoform groups based on their melting behavior, thereby enabling annotation-independent isoform detection ^22^. However, a systematic integration of long-read transcriptomics and global proteomics to directly match full-length transcript isoforms to experimentally detected protein isoforms in the same cell type has not yet been achieved.

Here, we present an integrated proteogenomic approach that combines SDS-PAGE-based protein separation with long-read transcriptome sequencing to systematically map mRNA and protein isoforms in the same cell type. By resolving proteins based on their intact molecular weight prior to mass spectrometry, we enhance the detection of distinct protein isoforms, including those generated by alternative splicing and proteolytic processing. While SDS-PAGE has historically been used to reduce sample complexity, its potential for protein isoform-level resolution is gaining renewed attention ^23,24^. Using this strategy in human RPE-1 cells, together with full-length transcriptome profiling, we establish a comprehensive and matched landscape of transcript and protein isoforms, revealing the molecular diversity encoded by the human genome. Our integrated proteogenomic isoform landscape indicates that a large fraction of alternative transcripts are translated, and highlights proteolytic processing as a major, underappreciated contributor to proteome complexity.

## Materials and Methods

### Cell Culture

hTERT RPE-1 cells were purchased from ATCC (CRL-4000) and maintained in DMEM/F-12 (1:1) + GlutaMAX^TM^ medium (GIBCO 31331-093) containing 10% fetal calf serum (FCS) (PAN-Biotech). Cells were cultured at 37°C in a humidified incubator of 5% CO_2_. Throughout the experiments the cells were kept under 15 passages.

### Sample Preparation

**Lysis**. hTERT RPE-1 cells were seeded on 15cm plates until ∼80% confluency, then washed and scraped with ice-cold PBS, collected into 15ml falcons, washed again and centrifuged (500g, 3min, 4°C). Cell pellets were lysed at room temperature (20mM Tris-HCl pH 7.5, 100mM NaCl, 1% SDS, 1mM MgCl2, Protease inhibitors) and immediately boiled (95°C, 5min). 1µl of Benzonase was added to digest DNA (37°C, 20min). Lysates were then centrifuged (14,000 rpm, 22°C, 15min) and the protein concentration in the supernatant was measured via BCA assay (Pierce BCA Protein Assay Kit). The proteins were then reduced (15mM DTT, RT, 30min), alkylated (20mM iodoacetamide, RT, 30min) and quenched (30mM DTT, RT, 30min). To desalt the protein lysate Acetone/Methanol precipitation was done. Namely, 6 volumes of -20°C Acetone/Methanol (8:1) was added to 1 volume of lysate, vortexed 20s and placed at -80°C overnight. Next, the samples were centrifuged (14,000rpm, 4°C, 30min), the supernatant was discarded and pellet was washed again with Acetone/Methanol and centrifuged (14,000rpm, 4°C, 10min). The pellet was dried and then solubilized in 2% SDS.

#### Protein Fractionation using GELFrEE® 8100 Fractionation system (Expedeon)

The proteins were fractionated according to their electrophoretic mobility employing the GELFrEE (Gel-Eluted Liquid Fraction Entrapment Electrophoresis) fractionation system with liquid phase recovery. We used the 5%- (mass range: 75-500kDa), 8%- (mass range: 35-150kDa) and 10%- (mass range: 15-100kDa) cartridges to generate 28 protein fractions (in total, 84 fractions per sample) according to the manufacturer instructions. Briefly, 150µl of protein sample (112µl-desalted protein, 8µl 1M DTT, 30µl Acetate Sample Buffer X5) was heated (95°C, 5min) and loaded onto the correspondent cartridge. The protein fractions were collected into 96 well-plates.

#### SP3-Digestion by BRAVO Automated Liquid Handling Platform (Agilent)

130µl of each protein fraction was transferred into a BRAVO-designated 96-well plate (28 fractions of each cartridge combined in one plate) along with two correspondent 96-well plates, one with magnetic beads (SeraMag beads A and B), and one with Trypsin-LysC (in 50mM Hepes pH 8). The digestion took place on the beads (1:25-50 protein:enzyme ratio).

#### TMT-labeling and HpH fractionation

We estimated the amount of peptides in the fractions to be 4µg to 8µg. The digested fractions were then labeled with TMTpro reagent set (Thermo Fisher Scientific, Cat #: A44520, Lot #: UL296296) according to the manufacturer’s instructions. Samples assignments: every 28 fraction set was divided into two separate plexes, where the “odd” fractions (fraction 1, fraction 3, .., fraction 27) were randomly assigned to one plex and the “even” fractions (fraction 2, fraction 4, .., fraction 28) where randomly assigned to a second plex. Assignments, plex #1: fraction 1 – 131N, fraction 3 – 127N, fraction 5 – 129N, fraction 7 – 132N, fraction 9 – 128C, fraction 11 – 127C, fraction 13 – 129C, fraction 15 – 131C, fraction 17 – 130C, fraction 19 - 126, fraction 21 – 130N, fraction 23 – 133C, fraction 25 – 132C, fraction 27 – 128N, Std1 - 133N, std2 - 134N. plex #2: fraction 2 – 129C, fraction 4 – 133C, fraction 6 – 127C, fraction 8 – 129N, fraction 10 – 132N, fraction 12 – 128N, fraction 14 – 126, fraction 16 – 130N, fraction 18 – 132C, fraction 20 – 130C, fraction 22 – 131C, fraction 24 – 128C, fraction 26 – 131N, fraction 28 – 127N, Std1 - 133N, std2 - 134N. Std1 and Std2 were whole cell lysates of hTERT-RPE1 cells that were not fractionated but went through the SP3-digestion (as described above). Std1 and Std2 contained 6µg and 60µg, respectively. The peptides from each plex were pooled, desalted with SepPak columns (manufacturer) and dried (speed-vac). We used the Dionex 3000 system (Thermo Fisher) to offline fractionate the desalted peptide mixture with high pH reverse phase fractionation (2p1mm C18 Xbridge Waters). Pooled peptides were resuspended in buffer A (0.0175% vol/vol NH4OH; 0.01125% vol/vol FA + 2% MeCN), loaded, and fractionated (96 min gradient from 0% B to 40% B in 70 min, 44% B in 4 min, 60% B in 5 min and kept at 60% B for the remaining 17 min; Buffer B: 0.0175% vol/vol NH4OH; 0.01125% vol/vol FA + 90% MeCN). Each 1 min a fraction was collected (in total 96 fractions). The automated collection procedure resulted in a total of 24 fractions, after recombination of selected fractions: (fraction i) + (fraction i + 24) + (fraction i + 48) + (fraction i + 72), i = 1 → 24. The fractions were dried and kept in -80°C.

#### MS Measurements

Samples were resuspended in Buffer A (3% acetonitrile, 0.1 formic acid). Peptides were separated with an easy-nLC System (Thermo Scientific) on a selfpacked column (20cm, 75µm ID, DrMaisch 1.9µm AQ) with a reverse phase gradient of increasing buffer B concentrations (90 % acetonitrile, 0.1 formic acid; 0 min 2 % B, 1min 4% B, 67 min 20% B; 20 min 30% B; 10 min 60% B; 1 min 90% B). Peptides were subsequently measured over ESI spray on an Q-Exactive HFx MS System (Thermo Scientific) for the 8er and 10er gels or on an Exploris480 MS System (Thermo Scientific) for the 5er gel. Settings on the HFx were: MS1 Res. 60k; AGC target 3e6; MaxIT 10ms; scan range 350-1500 m/z; MS2 Res 45k; AGC target 1e5; MaxIT 86ms; TopN 20; IW 0.7 m/z; Scan Range 350-1500 m/z; NCE 30, Dynamic Exclusion 30s.

Settings on the Exploris were: MS1 Res. 60k; normalised AGC target 300%; MaxIT 50ms; scan range 375-1500 m/z; MS2 Res 45k; normalised AGC target 100%; MaxIT 86ms; Cycle Time 1s; IW 0.4 m/z; Scan Range 350-1500 m/z; HCD Collision Energy 31%, Dynamic Exclusion 20s.

#### Generation of FLAG-tagged genes

TEAD3, NELFA, RRP1B and CTTNBP2NL genes were ordered in a pTwist_ENTR Kozak vector (Twist Bioscience). Using the Gateway Clonase II system (Thermo Fisher Scientific) the genes were cloned into the vectorspDEST_pcDNA5_BirA_FLAG_Nterm and pDEST_pcDNA5_BirA_FLAG_Cterm (Couzens et al, 2013). Sequencing validated the resulting products. Gene sequences were optimized, don’t contain introns and are therefore different from the human sequences.

#### Transfections

hTERT-RPE1 cells were seeded on 6-well plates and following the manufacturer’s instructions transfected with 5µg of the DNA constructs using Lipofectamine 3000 (Thermo Fisher). The cell lysates were prepared following the same procedure described in Sample preparation. The samples were run on 4-12% Bis-Tris gradient gels (NuPAGE, Invitrogen) and then blotted on PVDF membrane (Immobilon-P). Ectopic expression was detected using anti-DYKDDDDK Tag Antibody HRP Conjugate (2044, Cell Signaling).

### Next-generation sequencing (NGS) for total RNA

Total RNA from human retinal pigment epithelial-1(RPE-1) cells was extracted following the manufacturer’s protocol. Total RNA was extracted by TRIzol® Reagent (Life Technologies). 1 μg total RNA was used to prepare the VAHTS Stranded mRNA-seq Library Prep Kit (Vazyme).

After quality control, the libraries were sequenced in paired-end 2×150 nt manner on Illumina NovaSeq 6000 platform.

### PacBio library preparation and sequencing

PacBio library preparation and sequencing were performed as described before ^25^. Briefly, Total RNA from RPE-1 cells was extracted using TRIzol reagent (Life Technologies) following the manufacturer’s protocol, and then was converted to cDNA using the Clontech SMARTer PCR cDNA Synthesis kit. Then we proceed to PCR cycle optimization, and use the cycle number to generate a sufficient amount of double stranded cDNA products. After purification of PCR products by using AMPure purification methods, SMRTbellTM template preparation was performed. The libraries were sequenced on PacBio RSII SMRT platform according to the manufacturer’s instruction (Magbead mode for fragments >3 kb and Diffusion mode for all other fragment sizes; SMRT Cell v3 with 1200 min movie time). After sequencing, the raw reads were processed through the SMRT-Portal analysis suite (PacBio) to generate subread sequences for further analysis.

### Full-length transcriptome construction

#### Pre-processing

The primer of raw PacBio sequencing reads was removed using lima (version 2.0.0, --isoseq

--peek-guess). Then, the concatemer and poly(A) were removed by IsoSeq3 refine (version

3.4.0, --require-polya). For NGS sequencing, the clean reads were generated by fastp (version 0.21.0) with default parameters.

#### NGS-based error correction of PacBio sequencing reads

Proovread (verision 2.14.1, with default parameters) and LoRDEC (verision 0.9, with -k 17, -k 19 and -k 21, separately) were used to correct the sequencing errors in PacBio sequencing reads by taking advantage of the more accurate NGS reads. PacBio reads were corrected by Proovread and LoRDEC in parallel. After correction, GMAP (version 2021-02-22, Genome GRCh38, --sampling 1 -B 2 -n 0 -k 15 --min-intronlength 20) was used to align the corrected reads and reads with the best alignments (the longest alignment with the minimum number of mismatches) were kept for further analyses.

#### Clustering and collapsing

IsoSeq3 cluster (version 3.4.0) was applied to cluster and get the consensus sequences from the corrected PacBio sequencing reads. Pbmm2 align (version 1.9.0, with --preset ISOSEQ) was used to align the consensus sequences to genome GRCh38 and cDNA_Cupcake (collapse_isoforms_by_sam.py) was applied to collapse any identical isoforms based on the aligned exonic structure.

#### Annotating, quantification and filtering of full-length transcripts

SQANTI3 (version 3.6.1, with default parameters) was used to annotate the full-length transcripts with known GENCODE transcripts (gencode.v40.comprehensive.annotation.gtf downloaded from GENCODE) as well as discover novel transcripts. First, sqanti3_qc.py was used to annotate full-length transcripts by inputting transcript abundance information (parameter -e) and splicing junction information (parameter -c). The transcript abundances and splicing junction information was obtained from two NGS replicates, computed by kallisto quant (version 0.46.0, with default parameters) and STAR (version 2.7.8a, with --outFilterMultimapNmax 1).

Second, sqanti3_RulesFilter.py (-a 0.9) was used to remove potential artifacts caused by RT-template switching, intra-priming, etc. ORFs of the full-length transcripts were annotated by SQANTI at the same time.

#### Integration of full-length transcripts and ORFs with GENCODE annotation

Before integration, we first removed full-length transcripts (FLTs) with potentially degraded 5’end. For each FLT, we used SQANTI to compare its relationship with the annotated transcripts in GENCODE. For FLTs that were not annotated as “Full-splice match” by SQANTI, we checked if they were artifacts resulting from degradation at the 5’ end of the transcript due to technical limitations. Specifically, we compared the splice junction positions between FLTs in pairs; for all FLTs with the same “splice-junction-structure,” we retained only the longest transcript for further analysis. Also, for transcripts annotated in GENCODE, we used kallisto quant (version 0.46.0, with default parameters) to quantify each transcript and only kept transcript with mean TPM>1. Next, we integrated filtered FLTs from PacBio with filtered transcripts from GENCODE by splice-junction structure, and obtained the union set. For each transcript in the union set, we tried to set a label for its source (Fig. 4A). If the transcripts contain common splice-junction structure in both PacBio and GENCODE, they were labelled as “PB_&_GENCODE”, indicating the PacBio FLTs are totally “full-splice-match” compared to GENCODE transcripts; If the transcripts contain unique splice-junction structure in PacBio or GENCODE, they were labelled as “PB_only” or “GENCODE_only”. It should be mentioned that some FLTs are incomplete at 5’- or 3’- end likely due to technology limitation and they are partially “full-splice-match” compared to GENCODE transcripts. In this scenario, we hypothesized they are supported by both PacBio and GENCODE, and the PacBio FLTs should be elongated based on splice-junction structure of GENCODE transcripts. We set the label “GENCODE_from_PBcorrected” for this group of transcripts.

After the integrated transcript annotation was obtained, the ORF for each transcript can be inferred and the source of ORFs can be defined. Similarly, we classified all ORFs into four categories with the following priority: ’PB_&_GENCODE’ > ’GENCODE_from_PBcorrected’ > ’PB_only’ > ’GENCODE_only’. We also checked whether the ORF is from the canonical transcripts defined by Ensembl dataset and estimated the expression of ORF using the average TPM of all corresponding transcripts.

### Basic Data analysis

Raw Files from each Gel preparation were analysed separately with MaxQuant v2.0.3.0 ^26^ using the TMT module and search against the Uniprot database (UP000005640, 20220603) including listed chain variants of each protein sequence. Further data analysis was performed on the resulting peptides.txt table. The obtained 28 fractions from one gel separation were divided into two TMT experiments using 14 from the available 16 channels. Thus batch correction had to be applied by normalising each peptide’s intensities against the reference channel intensity if this peptide was quantified. In rare cases, when a peptide was not identified in the reference channel, peptide intensities were normalised against the average observed intensities in the reference channel in each TMT experiment. Next, peptides were z-scored and all sequences were remapped to Ensembl Gene IDs using Uniprot.ws and biomaRt R-packages.

Consequently, the subseeding analyses were performed in a gene centric manner.

### Peakfinder Analysis

Peptide traces from each gene were clustered using hdbscan from the dbscan R-package (1.1-10) using hdbscan function with the option minPts set to 2 and 3. For each clustering round noise was added to the data using the R function jitter on an initial value of -1 for each fraction and setting amount to -0.5. Resulting Clusters with more than 30 peptides were subjected to an additional clustering round using hdbscan. For each cluster the silhouette (cluster R-package 2.1.2) was calculated and the clustering event with the highest silhouette was picked for further analysis. After clustering with hdbscan, these noise traces were removed again from the data. Hdbscan itself reports noisy traces as cluster number zero. Peptide traces assigned to 0 were further considered to be noise and excluded from the subsequent peak detection. Next, the median profile of each cluster was calculated and peaks were determined using the findpeaks function (pracma 2.3.3 R-package) with the settings nups = 1, ndowns = 1 and threshold = 0.1.

Peak position was refined by determining the cross section of two linear models from both sides of the peak. Intensity and fraction confidence was determined in addition by running 20 bootstrap rounds with all peptides in a cluster.

Since different peaks of HDBSCAN clusters can represent the same protein isoform, we merged peaks with sufficiently similar molecular weights. In this way, peaks were either merged within each gel separately (gel centric analysis) or merged across all gels (gel combined analysis) using the following procedure: Briefly, we selected the peak with the highest z-score as the “seed” of an isofrac isoform. We then tested the overlap of adjacent peaks to the same isofrac isoform if they fall within the peak boundaries +/- 5000 Da. We therefore generated random distributions based on the Peak Apex and “start” and “end” of a peak. Then we sampled 20 data points for each peak and tested against the null hypothesis that the compared peaks are equal using the Wilcoxon Rank Sum test, as implemented in R. We did this in a bootstrap fashion, going for 50 rounds. Finally, we considered peaks to be similar if at least one resulting p-value is higher than 0.1.

To assess our ability to detect known protein isoforms, we compared isofrac isoforms with the Uniprot database, which includes information of splice variants and proteolytic cleavage variants. Specifically, we compared each isofrac isoform to all annotated Uniprot isoforms of the same gene. For each pair of isoforms, we computed the fraction of IsoFrac peptides shared with Uniprot (IsoFrac coverage), the fraction of Uniprot peptides shared with IsoFrac (Uniprot coverage) and the statistical significance of the peptide overlap (hypergeometric test p-value).

Since the ability to obtain significant p-values depends on the fraction of all peptides matching to a given uniprot isoform, we also calculated this fraction (total uniprot fraction). This results in a feature table for each IsoFrac Uniprot pair (with: IsoFrac coverage, UniProt coverage, hypergeometric test p-value, total uniprot fraction, all peptides identified per gene). The entire procedure was then repeated after randomly assigning peptides to IsoFrac isoforms. We then built a regression model using random forest (h2o package) using 10 fold cross validation (nfolds = 10). The obtained AUC from the ROC indicated that the prediction substantially improved with these scores in comparison of using each feature alone. Finally we used these scores to filter for the highest scoring isoform uniprot associations and calculated Δmass (Uniprot MW - Isoforac MW) and Quotient Mass (Uniprot MW/Isoforac MW) values to independently evaluate the filtering efficiency on the resulting mass difference between Uniprot and IsoFrac isoforms.

### Data availability

The raw and processed RNA-seq data generated in this study are deposited in the Gene Expression Omnibus under the accession number GSE261771 (It will be publicly available once the manuscript is accepted. Reviewers can access data through https://www.ncbi.nlm.nih.gov/geo/query/acc.cgi?acc=GSE261771 with token sdwzsayyjrathwb). The mass spectrometry proteomics data have been deposited to the ProteomeXchange Consortium via the pride partner repository^27,28^ with the dataset identifier PXD062269 (**Username:** **Password:** NkdkLWFiKCgB).

R-functions for processing the MaxQuant output tables are available under https://github.com/SelbachLab/IsoFrac.

The dataset can be interactively explored on the website proteinisoforms.mdc-berlin.de (username: “isofrac-review”, pw: “Ng0%rCK3jo%6@Wbs63”). Proteinisoform candidates as identified by IsoFrac are provided as a supplemental tab separated table (SupplementalTable 1).

## Results

### Peptide Correlation Profiling (PepCP) Resolves Protein-Level Information from Peptide Data

Correlating the abundance of proteins across biochemical fractions under non-denaturing conditions, so-called *protein correlation profiling* (PCP), is a well established method to map cellular proteins to organelles or protein complexes ^29^. For example, fractionating proteins via sucrose density gradient centrifugation, gel filtration or non-denaturing gel electrophoresis yields protein abundance profiles across fractions. The distribution of correlated proteins across these fractions then provides information about their subcellular localisation and/or multiprotein complexes they are members of ^30–37^.

Building on the principles of PCP, we developed *peptide correlation profiling* (PepCP) as a strategy to extract protein-level information from peptide-centric (bottom-up proteomic) data (Fig. 1A). PepCP follows four key steps: (1) Proteins are separated by SDS-PAGE based on their molecular weight into fractions ^38^. (2) Each fraction is enzymatically digested into peptides. (3) Peptides are identified and quantified across all fractions using quantitative mass spectrometry-based proteomics. (4) Peptide abundance profiles across fractions are analyzed to infer protein-level information.

**Figure 1:**
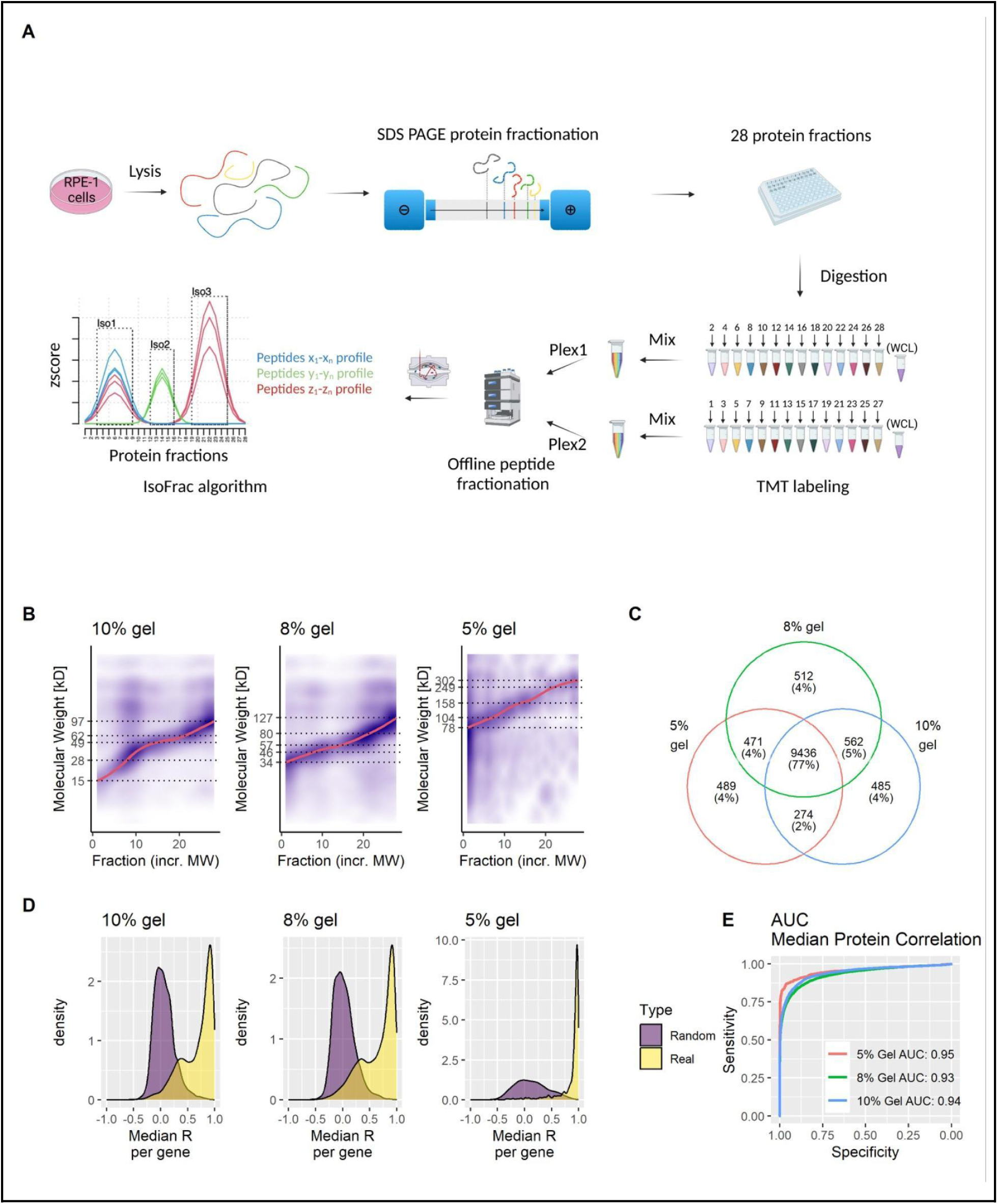
PepCP enables protein identifications from peptide data. **A)** PepCP sample preparation pipeline. **B)** Separation performance of different gels. Molecular weight projection among all collected fractions for the three gels used (10%, 8% and 5%). Protein MW are inferred from the major protein in each protein group for each identified peptide based on the highest z-score across all fractions per gel. The loess regression trendline is depicted for each gel. **C)** Venn diagram of all identified GenCode ID Groups among the different gels. **D)** Distribution of median average Pearson's coefficients of correlation between all peptide pairs within a gene (Real) or random pairs between peptides of a gene and peptide from other genes (Random). **E)** Receiver operating characteristics from the corresponding distributions in D). AUC is depicted in the figure.

We extracted proteins from the diploid human retinal pigmented epithelial cell line RPE-1 and fractionated them by SDS-PAGE using the GELFREE system, which improves protein recovery ^39^. To cover isoforms across molecular weight ranges we used gels with 10%, 8% and 5% acrylamide. A total of 28 fractions per gel were collected, and proteins were digested using an automated SP3 workflow ^40^. Peptides were labeled with tandem mass tags (TMT) to enable quantification across fractions ^41^. To achieve this, the 28 fractions were combined into two TMT-16 plexes, each containing 14 fractions, along with an unfractionated whole-cell lysate (WCL) reference channel. This design allowed peptide abundance profiles across SDS-PAGE fractions to be quantified as ratios relative to the reference channel. Combined peptide mixtures were separated offline by high-pH reverse-phase HPLC and subsequently analyzed by LC-MS/MS.

We processed the data using MaxQuant with a protein database that integrated UniProt sequences and isoforms arising from proteolytic cleavage events reported in UniProt (see Methods). In total, we identified ∼150,000 peptides (FDR 1%) in ∼13,000 protein groups that we mapped to ∼12,000 genes (Fig. 1C). For downstream analysis, we only considered peptides that uniquely mapped to a single gene. After remapping all sequences to human Ensembl gene annotations and filtering for genotypic peptides, we retained 136,752 peptides in 12,176 protein groups, corresponding to 9,288 genes.

To assess the resolving power of our SDS-PAGE gels, we extracted the annotated molecular weights of all unique peptides in each fraction and visualized their distribution as a density plot (Fig. 1B). As expected, protein molecular weight increased with fraction number, spanning a range of 15 to 250 kDa across all three gel types. Consistent with their increasing pore sizes, the mass ranges captured by the 10%, 8%, and 5% gels also increased accordingly.

The premise of Peptide Correlation Profiling (PepCP) is that peptides derived from the same protein isoform should exhibit correlated abundance profiles across fractions. If each gene encoded only a single protein isoform, all corresponding peptides would be expected to follow similar abundance profiles. To assess this globally, we mapped peptides to the genome (see Methods) and computed pairwise correlations for all peptides mapping to the same gene. The vast majority of peptide pairs showed positive correlations (r > 0), with a major peak observed around r = 0.9 (Fig. 1D). This confirms that peptides mapping to the same gene tend to have correlated abundance profiles across protein fractions. As expected, correlations between randomly sampled peptide pairs were distributed around r = 0. To further evaluate the performance of PepCP, we computed receiver operating characteristic (ROC) curves (Fig. 1E). Both specificity and sensitivity were high, particularly for the 8% and 10% gels, which achieved area under the ROC curve (AUC) values of 0.93 and 0.94, respectively. Notably, these estimates are conservative, as many genes encode multiple protein isoforms, which is expected to lower peptide-peptide correlations.

Having demonstrated that PepCP can reliably assign peptides to proteins, we next investigated whether it could provide insights into protein isoforms. As a first test, we analyzed intronless genes, which are expected to encode a single protein isoform ^42^. Consistent with this expectation, we observed a single peak for this gene category (Fig. 2A).

**Figure 2:**
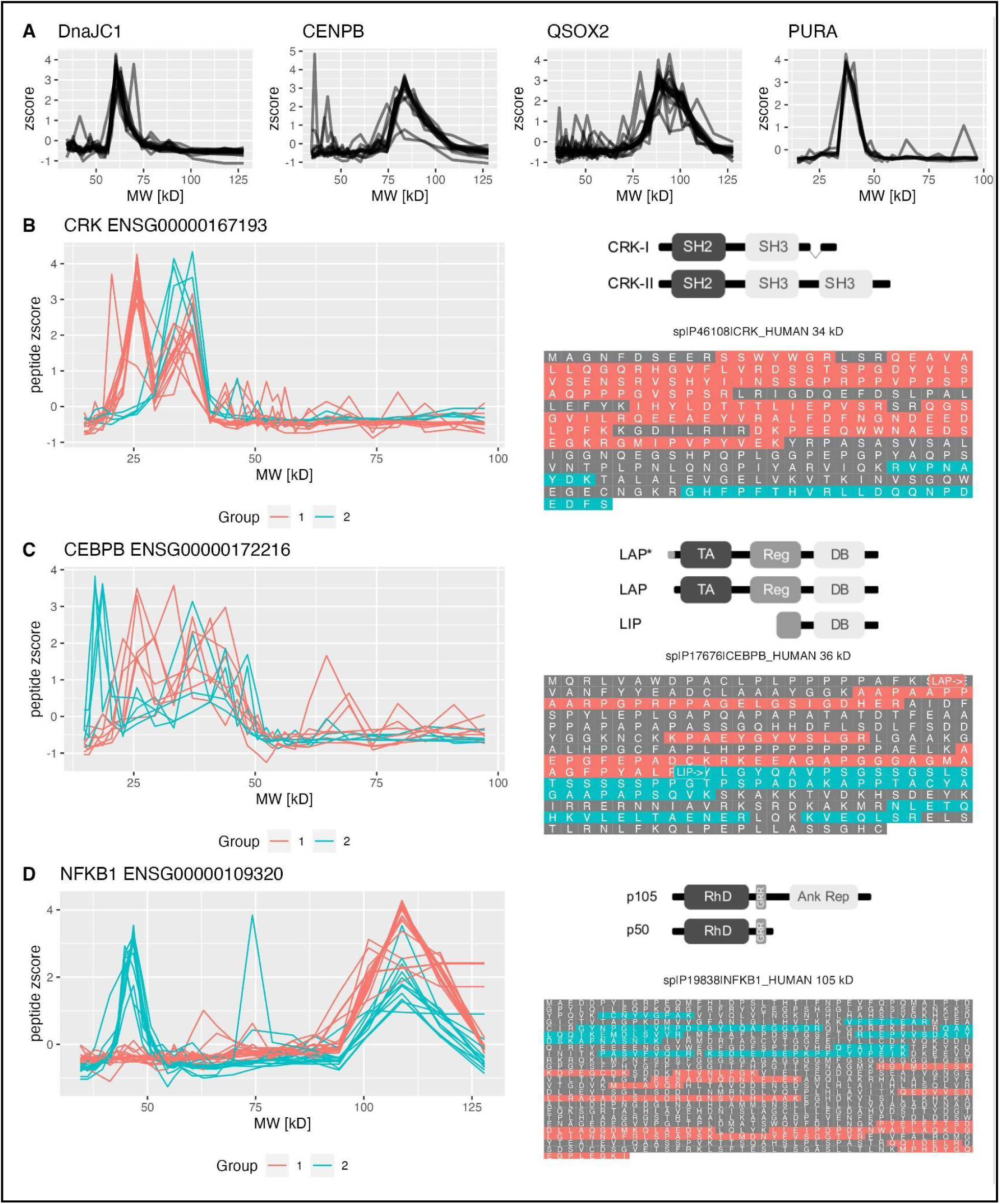
Example peptide separation profiles. Peptide zscore traces across SDS gel fractions for selected intronless genes **(A)**. **B-D** show observed protein isoform traces generated by different processes: Alternative splicing **(B)**, alternative initiation **(C)** and protein processing **(D)**. For all traces fraction numbers have been converted to MW values based on average MW observed among identified proteins in each fraction. Identified peptides were mapped to known protein sequences for each corresponding gene. The position of identified peptides across each canonical protein sequence is depicted on the right side (B-D). Peptide-groups were assigned manually.

In principle, protein isoforms can arise through three major mechanisms: (i) alternative splicing ^5^, (ii) alternative translation initiation or termination ^43,44^ and (iii) posttranslational processing, such as proteolytic cleavage ^45^. Importantly, we found that PepCP can provide information on all of these processes:

i. Alternative splicing: We identified both splice isoforms of CRK, where peptides unique to the larger isoform exhibited a single peak, whereas peptides shared by both isoforms showed two distinct peaks (Fig. 2B).
ii. Alternative translation: We observed isoforms of CEBPB that arise from alternative translation initiation (Fig. 2C).
iii. Proteolytic processing: We identified the processed isoform of NFKB1, formed by protein cleavage (Fig. 2D).

These findings illustrate that PepCP enables the detection of distinct protein isoforms, providing a more comprehensive view of proteome complexity.

### Mapping the Protein Isoform Landscape

The examples highlighted above demonstrate PepCP’s ability to provide detailed information about protein isoforms. To globally analyze these data, we developed IsoFrac, a computational pipeline that automatically assigns peptides from PepCP data to protein isoforms (see Methods). Briefly, the IsoFrac algorithm consists of three steps (Fig. 3A):

1. Peptide clustering: Peptide abundance profiles are clustered to identify groups of peptides with similar fractionation patterns. We used hierarchical density-based spatial clustering of applications with noise (HDBSCAN), an advanced algorithm that optimizes cluster stability while excluding noisy data ^46^.
2. Peak detection: For each peptide cluster, an average abundance profile is computed, and peaks are identified.
3. Isoform definition: Isoforms are assigned based on peak positions. Peptides with peaks in close proximity are merged into the same isoform.

**Figure 3:**
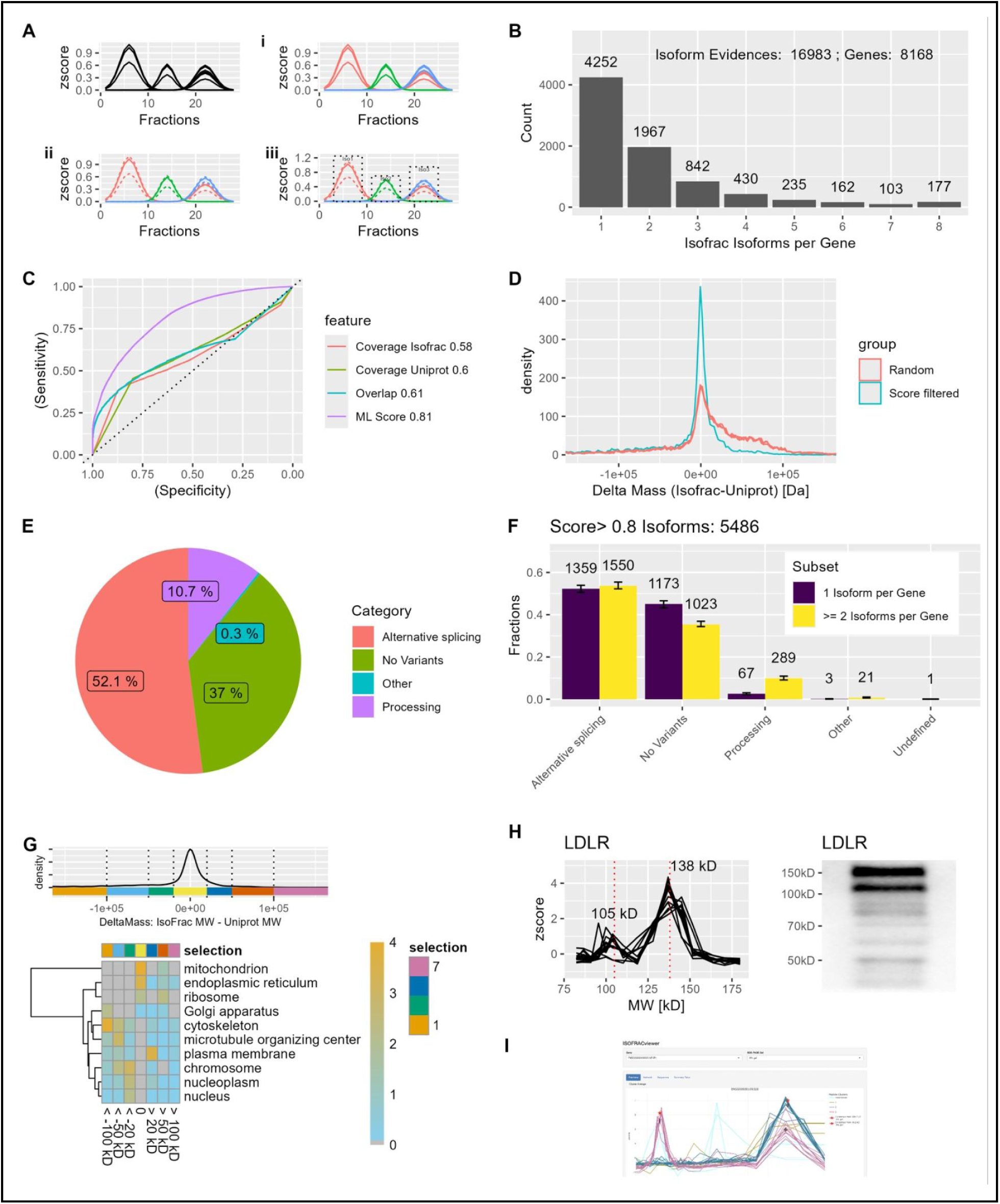
IsoFrac identifies protein isoforms on a global scale. **A)** Diagram illustrating IsoFrac Peak detection. (i) All peptide traces are clustered using hdbscan. (ii) Average trace for each cluster is calculated. (iii) Peaks are identified based on each average cluster trace and combined across clusters and gels. **B)** Number of identified isoforms per gene. **C)** ROC curves for single metrics and combined regression score to assess the quality of isofrac uniprot pairs. **D)** Δmass density curves for the different subsets ‘Score filtered’ (selection for highest Score) and ‘Random’ (random assignment). **E)** Fractions of assigned protein variant categories for a high confidence set (Score > 0.8). **F)** Comparison of protein fractions from subfigure E for proteins with >=2 isoforms in Uniprot (purple) against proteins with a single entry (yellow) in Uniprot. **G)** Enrichment analysis of proteins from different bins among the Δmass distribution of IsoFrac vs Uniprot protein molecular weights from proteins with high confidence assignments (score > 0.8). −log10 pvalues are projected as a heatmap for cellular component gene ontologies with the different bins from Δmass distribution shown in separate columns. **H)** LDLR (95 kD) peptide traces in the 5% gel with two peaks at ∼105 kD (unmodified) and ∼138 kD (left). The western blot result is depicted on the right side. **I)** Screenshot of ISOFRACviewer, an online tool to assess isoform information from this study.

Initially, IsoFrac defines isoforms separately for each gel (10%, 8% and 5%). These are then merged across gels if they exhibited similar molecular weights. Using this procedure, we identified a total of 16,983 isoforms derived from 8,168 genes (Fig. 3B). For roughly half of these genes we detected more than one protein isoform.

We next assessed whether IsoFrac reflects known protein isoform annotations. To this end, we compared each IsoFrac isoform to all annotated UniProt isoforms for the same gene using three features: (i) IsoFrac coverage – the percentage of IsoFrac peptides that match a given UniProt isoform, (ii) UniProt coverage – the fraction of UniProt peptides covered by IsoFrac peptides, (iii) Overlap – a hypergeometric test to assess the significance of the overlap between UniProt and IsoFrac peptides. As a negative control, we randomly assigned peptides from the same gene to IsoFrac isoforms and computed receiver operating characteristic (ROC) curves (Fig. 3C). As expected, all three features contributed useful information but with limited predictive power. We therefore trained a gradient-boosted classification tree model to integrate the three features into a single score (see Methods). This markedly enhanced predictive accuracy, achieving an area under the ROC curve (AUC) of 0.81 (Fig. 3C). Notably, since UniProt annotations do not fully capture the true diversity of protein isoforms, we do not expect an AUC of 1 even if our data were perfectly accurate.

To further validate our data, we tentatively assigned each IsoFrac isoform to the best-matching UniProt isoform. We then calculated the molecular weight difference (Δ mass) between the assigned IsoFrac and UniProt isoforms. We observed that these Δ masses were significantly smaller than Δ masses for randomly assigned UniProt isoforms (Fig. 3D).

Importantly, this provides independent validation of our data, as molecular weight information was not used during the isoform assignment process.

To assess global trends in our dataset, we assigned each IsoFrac isoform to its best-matching UniProt candidate and categorized it by its known isoform-generating mechanism. In addition, we only included isoforms with a good sequence identification score (ML_Score > 0.8, 5486 proteins). Based on these assignments, the majority of protein isoforms arise from alternative splicing, while approximately 10% are generated by posttranslational processing events (Fig. 3E).

Next, we grouped genes into two categories: those with a single protein isoform detected in IsoFrac and those with multiple isoforms (Fig. 3F). As expected, genes with more than one annotated isoform in UniProt were more likely to be identified with more than one isoforms in IsoFrac, whereas proteins with a single UniProt entry (“No Variants”) were more often detected as a single isoform. Surprisingly, however, we detected multiple protein isoforms for 1,023 genes annotated with only a single UniProt isoform, suggesting the presence of previously unrecognized protein isoforms.

To explore functional trends, we performed gene ontology (GO) enrichment analysis across different bins of the Δ mass distribution. The resulting heatmap of enrichment p-values revealed a Δ mass-dependent distribution of cellular protein localization (Fig. 3G). Since posttranslational modifications (PTMs) influence protein molecular weight, we expected larger modifications — such as glycosylation — to lead to an upward shift in observed molecular weight. Consistently, we found endoplasmic reticulum and plasma membrane proteins – both known to be frequently glycosylated – to be enriched in higher Δ mass bins. For example, LDLR (the low-density lipoprotein receptor), a glycoprotein, displayed two peaks in IsoFrac with an identical set of identified peptides (Fig. 3H, left). Western blot analysis confirmed this observation, showing two distinct bands in the corresponding molecular weight range (Fig. 3H, right).

Our IsoFrac dataset offers a comprehensive resource for global and gene-specific isoform analysis. To facilitate exploration, we developed an interactive online tool (Fig. 3I) that enables users to efficiently browse isoform profiles. The dataset is accessible at https://proteinisoforms.mdc-berlin.de (username: “isofrac-review”, pw: “Ng0%rCK3jo%6@Wbs63”).

### Long-Read Sequencing Connects Transcriptome Complexity to Protein Isoform Diversity

To complement our proteomic isoform detection and to consider potential RPE-1-specific alternative splicing, we systematically characterized the RPE-1 transcriptome using long-read sequencing. To achieve this, we combined full length transcriptome (PacBio) and short-read (NGS) RNA sequencing into a unified RNA isoform identification pipeline (Supplemental Figure 1). After integrating our data with transcript annotation in GENCODE (Fig. 4A), we identified 45,223 protein-coding transcripts with 32,612 open reading frames (ORFs) across 11,605 genes (Fig. 4B). Among these, 20,420 (45%) transcripts were detected in both our PacBio data and GENCODE, highlighting a strong concordance between full-length transcripts and GENCODE-annotated expressed transcripts. Additionally, 15,453 GENCODE-only transcripts lacked evidence in the PacBio dataset but were retained for further analysis due to their detection in our NGS data. Interestingly, 7,652 transcripts (17%) were detected exclusively by PacBio but not in GENCODE, potentially representing novel alternative splicing events. As an example, we examined the TMPO gene and found that our integrated annotation provides a comprehensive and non-redundant representation of RPE-1 transcripts and ORFs. Specifically, we identified seven transcripts and four ORFs. These correspond to the four major TMPO isoforms (Fig. 4C) and have also been identified on protein level by our IsoFrac pipeline (Fig. 4D).

**Figure 4:**
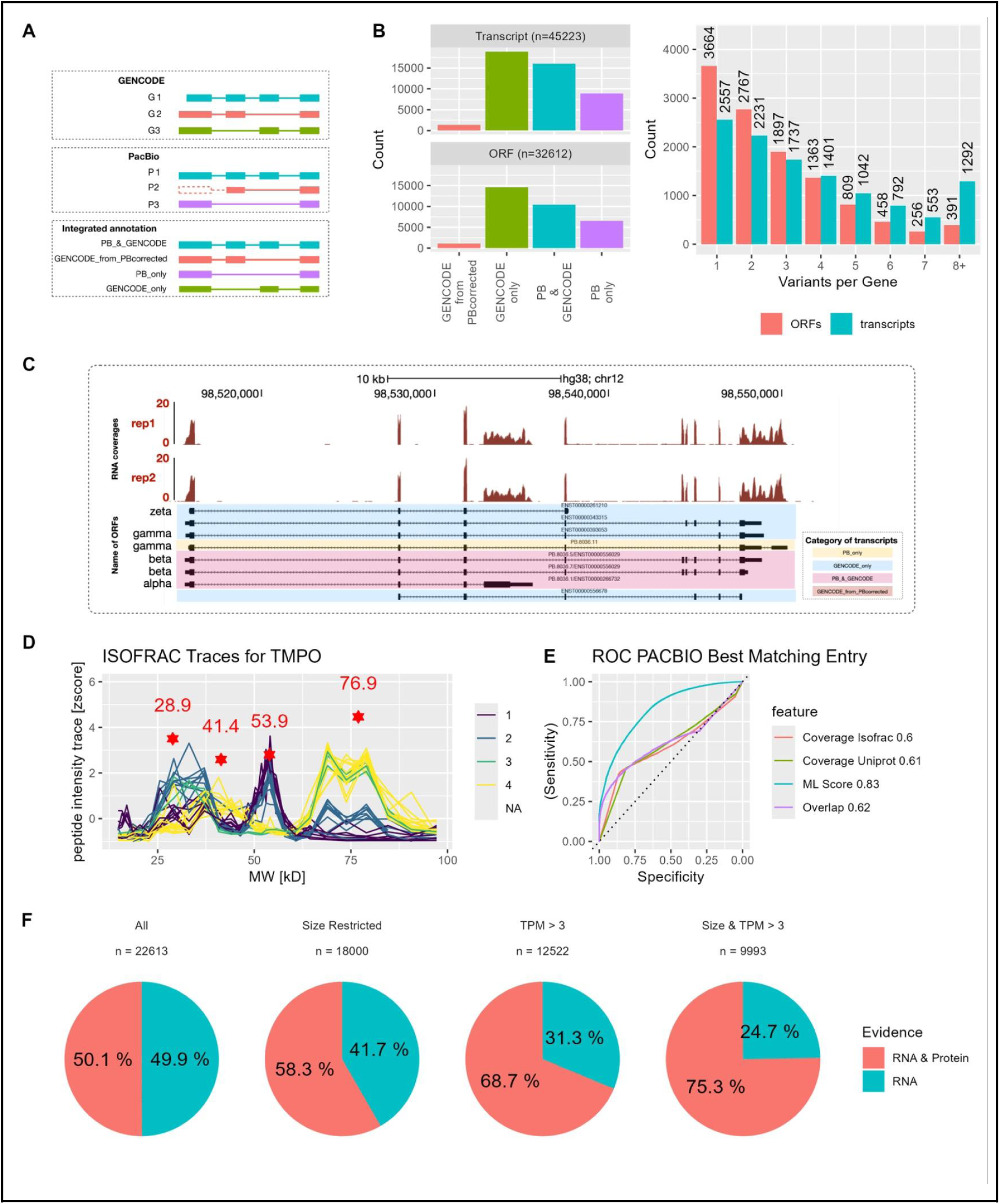
Integration of RNA PACBIO Sequencing data. **A)** Label assignment strategy of obtained transcripts. The upper block: transcripts from GENCODE annotation; The middle block: transcripts from PacBio; The bottom block: integrated annotation. See Method for details. **B)** Number of transcripts and ORFs per gene across different annotation categories (left) and across identified variants per gene (right). **C)** Visualization of RNA-seq coverage and isoforms for TMPO gene. Tracks in red: RNA-seq coverages from NGS, with two replicates; Blocks in black: transcripts in integrated annotation for TMPO genes, categories of each transcript are labelled with different colours. Names of ORFs are labelled on the left. **D)** Isofrac peptide traces across fractions for TMPO. Identified protein isoforms are depicted by a red star and the corresponding estimated MW in kD. **E)** ROC of different metrics for predicting the best matching PACBIO entry for each IsoFrac isoform. **F)** Percentage of isoforms detected at the RNA (green) or RNA and protein level (red). Different technical biases were considered in addition to have a more realistic estimate: 1. MW resolution of Isofrac: Proteins in the RNA database from the same gene were merged, if their MW difference is below 5 kD. 2. RNA abundance, proteins were excluded from the analysis, if the transcript abundance was below 3 TPM.

Next, we integrated protein isoform data with transcript isoforms by repeating our isoform mapping approach, this time using a custom protein sequence database derived from RPE-1-specific ORFs (PacBio database). To ensure compatibility with our previous analysis, we applied the same regression model used for UniProt-based isoform assignments. Using the PacBio database, we achieved a prediction performance of 0.83 (AUC), slightly surpassing the 0.81 AUC obtained with UniProt (Fig. 4E).

Having an integrated transcriptome and proteome isoform dataset allows us to directly assess the ongoing question of how much alternative splicing contributes to proteome complexity, a topic of active discussion and differing perspectives ^5,9^. To address this, we assigned each protein isoform to its best-matching ORF in an exclusive manner, meaning that once an ORF was assigned to a protein isoform, it was not reused for another assignment.

Using this approach, we mapped protein isoforms to approximately 50% of the 24,376 ORFs identified from genes detected in both the transcriptomic and proteomic datasets (Fig. 4F, left). At first glance, this might suggest that only a fraction of alternatively spliced transcripts are translated into stable protein products. However, a direct comparison between transcriptomic and proteomic datasets must account for technical differences between the two methods.

Specifically, our SDS-PAGE-based protein separation cannot distinguish protein isoforms with similar molecular weights, which may lead to an underestimation of protein diversity. To correct for this, we collapsed ORFs encoding proteins that differed by less than 10 kDa into a single entry, which increased the coverage to over 60% (Fig. 4F, Supplemental Fig. 2). Another factor is protein abundance, as low-abundant protein isoforms may escape detection. Indeed, when restricting the analysis to transcripts with TPM >3, coverage increased to over 70% (Fig. 4F, Supplemental Fig. 2). Combining both adjustments — collapsing ORFs by molecular weight and applying an abundance filter — further increased the protein detection rate to 78%. In summary, these findings indicate that most alternatively spliced transcripts that are detectable with our technology are indeed translated into stable proteins, supporting previous observations using different proteomic approaches ^21^.

### Most database specific protein assignments are protein cleavage products

Our dataset provides evidence for previously unannotated protein isoforms, as several genes exhibit multiple protein isoforms despite being annotated with only a single one (Fig. 3F). These isoforms could result from RPE-1-specific alternative splicing. To investigate this, we mapped each protein isoform to both UniProt and the PacBio database. More than 75% of isoforms showed identical assignment scores, indicating they mapped equally well to both references (Fig. 5A). Only a few hundred isoforms aligned more strongly with PacBio, suggesting that RPE-1-specific alternative splicing contributes to a relatively small fraction of the observed protein isoforms. This implies that most alternatively spliced mRNAs are already annotated in UniProt.

**Figure 5:**
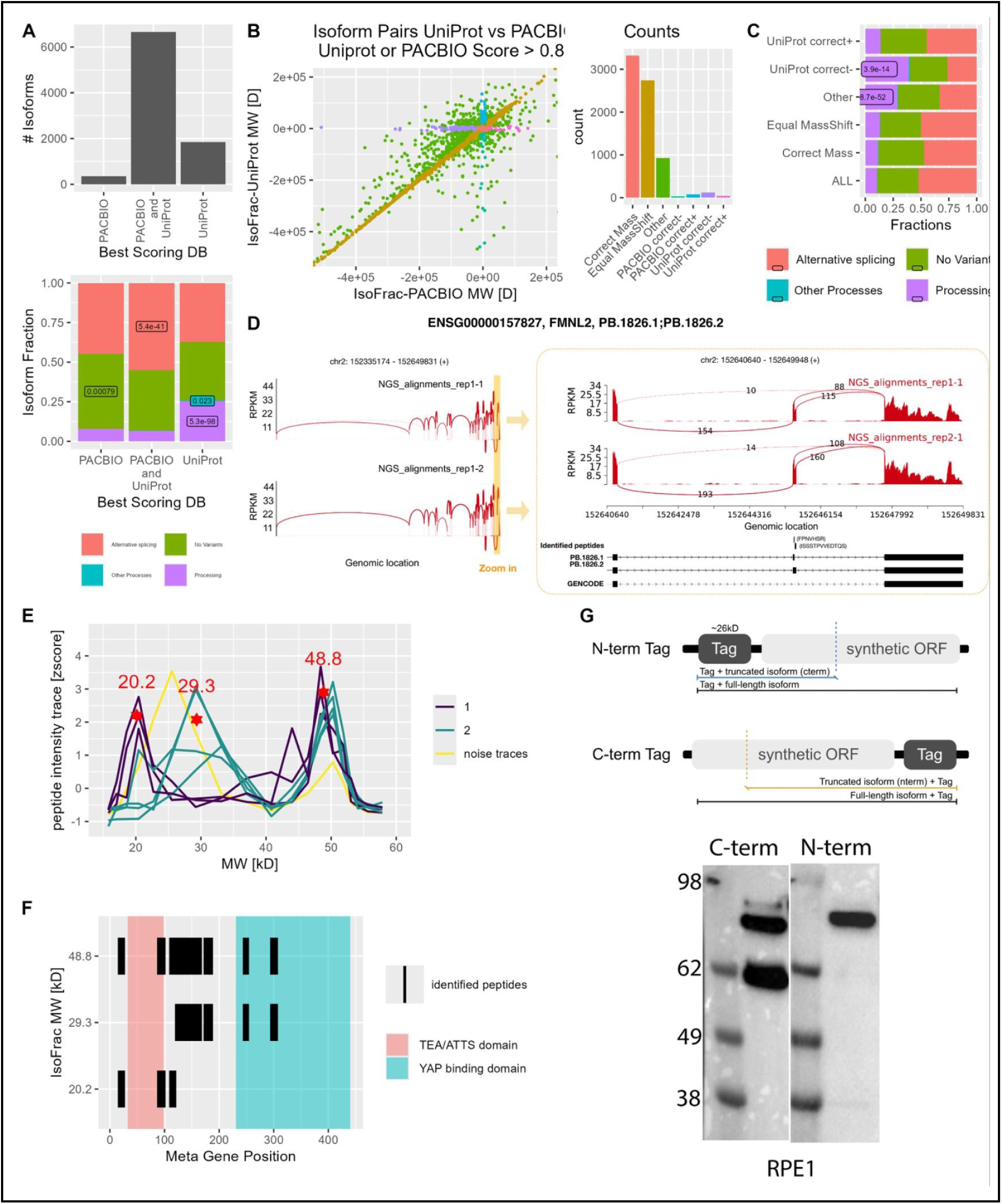
Validation of new protein isoforms. **A)** Number of identified Isoforms based on the best scoring candidate from PACBIO or UniProt (left barplot) and corresponding fractions of protein isoform generating processes (right barplot). P-values < 0.01 from an hypergeometric enrichment test are depicted in the figure. **B)** Pairwise Δmass comparison (IsoFrac Mass - Database Mass) of PACBIO against Uniprot matches for a selected subset (any score of an isofrac database pair > 0.8). Seven different groups have been defined, based on the position with this plot: “Correct Mass” (PACBIO and UniProt masses are similar to identified IsoFrac mass); “Same Offset Mass” (PACBIO and UniProt masses are similar but different from the corresponding IsoFrac mass); “PACBIO Zero +/-” (PACBIO mass is correctly assigned, but UniProt is +/- off); “UniProt Zero +/-” (UniProt mass is correctly assigned, but PACBIO is +/- off) and “Other”. The corresponding barplot depicts the numbers of isoforms belonging to these groups. Only Peaks with a Score > 0.8 in UniProt or PACBIO have been selected for the analysis. **C)** Enrichment analysis of Uniprot related Isoform categories within different Δmass categories from **(B)**. The p-value from hypergeometric testing is depicted in the figure for significantly enriched fractions. **D)** The Sashimi Plot demonstrates that short-read NGS, PacBio sequencing, and peptide alignments support the newly identified transcript of FMNL2 (ENSG00000157827). The left panel shows read coverage and splice junctions from NGS, while the right panel highlights peptide alignment to the transcriptome, including the novel transcript from PacBio and known GENCODE annotations. Arc labels indicate splice junction read counts. **E)** Peptide zscore traces from the Isofrac analysis. The three identified isoform candidates and their estimated MWs are marked with a red star in the figure. The western blot of TEAD3 is depicted on the right side. **F)** Meta Gene Position analysis of the peptides identifying the three different isoform candidates of TEAD3. **G)** Western Blot analysis of a TEAD3 pull down using a construct with either n-terminal or c-terminal Tag.

In contrast, many protein isoforms mapped better to UniProt than to the PacBio-derived database. To understand this discrepancy, we examined UniProt annotations and found that isoforms aligning more closely with UniProt were significantly enriched in the “processing” category. This aligns with expectations, as the PacBio database does not capture protein isoforms generated by posttranslational modifications.

To explore this further, we compared the observed molecular weights of protein isoforms to their best-matching entries in UniProt and PacBio. For over 3,000 isoforms, the observed molecular weight closely matched (within 10 kDa) annotations in both databases, supporting the accuracy of our approach. However, we identified cases where the molecular weight was consistent with UniProt but lower than the corresponding PacBio entry (“UniProt correct minus”). These isoforms were enriched in the “processing” category, further supporting the role of proteolytic processing in generating protein isoforms. For most isoforms with mass deviations, the shift relative to both UniProt and PacBio was similar, reinforcing the idea that alternative splicing alone does not explain most unannotated isoforms and highlighting the role of translational and posttranslational processing in proteome diversity.

While alternative splicing accounts for only a minor fraction of newly identified protein isoforms, we examined whether our proteomic data provide peptide-level evidence for novel splice variants detected in the PacBio dataset. We identified seven peptides that uniquely mapped to the PacBio transcriptome but were absent from UniProt at the time of analysis.

These peptides were confidently identified with scores comparable to annotated peptides (Supplemental Figure 3) and correspond to novel splice junctions (Fig. 5D and Supplemental Figures 4). One of the identified variants is generated by an alternative translation start site that was not annotated in the UniProt release used in our search (see Methods). Notably, this variant is now included in a more recent UniProt release, supporting the validity of our finding and demonstrating that our integrated long-read transcriptome and proteome analysis can anticipate updates to genome annotations (Supplemental Figure 4.5). Altogether, while alternative splicing is not the primary driver of novel protein isoforms, our approach successfully uncovers a subset of previously unannotated, alternatively spliced protein variants.

Given that many novel isoforms exhibit lower molecular weights than their best-matching UniProt entries, we hypothesized that they arise from posttranslational processing. For instance, we detected three isoforms of TEAD3, a transcription factor and key Hippo pathway effector ^47^. The full-length TEAD3 (48.7 kDa) contains an N-terminal DNA-binding domain and a C-terminal YAP-binding region. While SwissProt lists only this isoform, TrEMBL predicts a shorter N-terminal fragment (20.2 kDa). Our IsoFrac analysis detected both isoforms and revealed an additional C-terminal isoform (∼30 kDa) that lacks the DNA-binding domain (Fig. 5E, F). Since this isoform is absent from the PacBio data, we hypothesized that it arises from posttranslational processing. To test this, we generated N- and C-terminally tagged TEAD3 constructs and transiently expressed them in RPE-1 cells. To rule out alternative splicing as a source of different isoforms, we used codon-optimized cDNA lacking introns and differing in nucleotide sequence from the endogenous TEAD3 mRNA. Western blotting of the C-terminally tagged construct detected prominent bands for both the full-length and the additional C-terminal isoform, validating our findings (Fig. 5G). We also created N- and C-terminal tags for three additional proteins and observed bands consistent with PepCP-based isoforms (Supplemental Figure 5). In summary, our integrated proteomic and transcriptomic analysis suggests that while alternative splicing contributes to proteome complexity, posttranslational processing plays a more prominent role than previously appreciated.

## Discussion

Despite extensive efforts to characterize transcript diversity, a systematic mapping of protein isoforms has remained elusive. Here, we demonstrate that peptide correlation profiling (PepCP) enables the global identification of over 16,000 protein isoforms, providing a comprehensive view of proteome diversity in RPE-1 cells. By integrating these data with long-read transcriptomics, we provide an integrated landscape of mRNA and protein isoforms (https://www.proteinisoforms.mdc-berlin.de). Our data reveals that the majority of alternatively spliced isoforms that are technically detectable at the protein level are captured in the proteomic data. Notably, we also identify numerous protein isoforms that have not yet been annotated, with the majority likely arising from proteolytic processing rather than alternative splicing. Collectively, these findings highlight posttranslational processing as an important contributor to proteome complexity and underscore the importance of integrating transcriptomic and proteomic approaches for a more complete understanding of isoform diversity.

The systematic identification of protein isoforms remains a major challenge. In conventional bottom-up workflows, proteins are enzymatically digested into peptides, thereby severing the link between peptides and their protein isoform of origin—a limitation commonly referred to as the protein inference problem ^15^. One strategy to mitigate this issue is ultradeep bottom-up proteomics, which combines multiple proteases, extensive peptide fractionation, and diverse fragmentation methods to maximize sequence coverage ^21^. While this approach increases the likelihood of detecting isoform-specific peptides, it does not fundamentally resolve the ambiguity introduced by shared peptide sequences. Top-down proteomics, in contrast, analyzes intact proteins and can directly resolve protein isoforms ^11^. Recent advances have substantially improved its sensitivity and resolution, with applications now extending to single-cell analysis ^6,12,13,48^. However, top-down approaches still fall short of the proteome-wide coverage achieved by transcriptomics or deep bottom-up workflows. Our peptide correlation profiling (PepCP) approach circumvents these limitations by leveraging protein-level separation through SDS-PAGE prior to bottom-up mass spectrometry. This strategy directly resolves intact proteins by molecular weight and retains this information at the peptide level, allowing protein isoforms to be distinguished based on their migration profiles. Prior methods infer proteoform identity from co-variation of peptide abundance across biological perturbations or protein complexes ^18,19^, or from peptide melting behavior across cell lines ^22^. While such approaches offer valuable functional insights, they do not provide direct information on intact protein size or enable integration with full-length transcript isoforms from the same sample. Our SDS-PAGE-based approach enables the detection of protein isoforms even in the absence of unique peptides and provides orthogonal information on their apparent molecular mass. By integrating the resolving power of SDS-PAGE with the depth of shotgun proteomics, PepCP offers a scalable and broadly accessible strategy for proteome-wide identification of protein isoforms, helping to close the gap between transcriptome complexity and protein-level evidence.

While PepCP enables proteome-wide isoform detection, several limitations should be noted. First, the resolving power of SDS-PAGE is inherently restricted, making it difficult to distinguish isoforms with similar molecular weight. Second, for proteins with complex peptide migration profiles — due to broad peaks, overlapping species, or low signal-to-noise ratios — automated isoform assignment can be ambiguous. Finally, although SDS-PAGE primarily separates proteins by size, migration behavior can also be affected by charge, hydrophobicity, or residual secondary structure, potentially leading to inaccuracies in apparent molecular weight estimates ^49,50^. These factors highlight the need for cautious interpretation and suggest that integrating orthogonal separation strategies may further enhance isoform resolution in future studies.

The limited detection of isoforms in standard proteomic datasets has led to the view that many alternatively spliced transcripts may not yield stable protein products, and that most genes predominantly express a single protein isoform ^9,10^. However, by accounting for the technical constraints of our approach — particularly molecular weight resolution and protein abundance — we find that the majority of alternatively spliced transcripts do give rise to detectable protein isoforms. This finding is consistent with recent ultradeep shotgun proteomic studies and reinforces the notion that alternative splicing is a major contributor to proteome complexity ^21^.

Because our approach provides a global and annotation-independent view of protein isoforms, it enables an unbiased comparison between observed and annotated isoform diversity. According to existing annotations, most protein isoforms detected arise from alternative splicing. However, we also observe multiple isoforms for many genes that are annotated to produce only a single protein isoform, indicating that current reference annotations remain incomplete. Interestingly, alternative splicing appears to account for only a small fraction of this unexplained diversity: while our combined long-read transcriptome and proteome data identify several RPE-1–specific isoforms, these represent only a minor subset of the newly observed protein isoforms.

Multiple lines of evidence from our data suggest that proteolytic processing contributes more substantially to proteome diversity than previously appreciated. We detect a large number of protein isoforms that cannot be explained by alternatively spliced transcripts present in RPE-1 cells, and notably, most of these novel isoforms are shorter than their annotated counterparts.

Given our ability to efficiently detect proteolytic cleavage products—including many known processing events—these observations strongly suggest that a significant fraction of the newly observed isoforms arise through posttranslational proteolysis. While we cannot globally exclude alternative translation initiation or termination as potential sources of N- or C-terminal isoforms, this mechanism is unlikely to account for the isoforms we selected for validation. In these cases, including TEAD3, isoforms were detected following transfection of codon-optimized cDNAs with heterologous 5′ and 3′ UTRs, which are not expected to support alternative translation in the same way as endogenous transcripts. Importantly, it is also unlikely that these additional isoforms result from nonspecific degradation: protease inhibitors were included throughout the workflow, and the observed isoforms display well-defined molecular weights rather than the diffuse profiles expected from random cleavage products. More broadly, current isoform annotations may be inherently biased toward transcript-derived variants, as splicing isoforms can be readily inferred from RNA sequencing, whereas proteolytic isoforms require direct protein-level evidence for detection. As a result, a large fraction of the protein isoform landscape — particularly those shaped by posttranslational mechanisms — may remain uncharted.

Together, our findings provide a comprehensive and experimentally grounded view of isoform diversity at both the transcript and protein levels. By combining deep full-length transcriptome sequencing with peptide correlation profiling, we show that a substantial fraction of transcript isoforms are translated, and reveal that posttranslational processing contributes significantly to the generation of protein diversity—often beyond what is captured by current annotations. These insights not only highlight the importance of integrated proteogenomic approaches for isoform discovery, but also underscore the need to revise existing models of proteome complexity that focus predominantly on alternative splicing. As methods for intact protein analysis continue to advance, and as more cell types and conditions are explored, we anticipate that similar strategies will enable increasingly detailed and dynamic maps of the human protein isoform landscape.

## Supporting information

Supplemental Table 1 Column Descriptions

## Acknowledgements

We thank Christian Sommer and Mohamad Haji for excellent technical assistance, and Christopher Buccitelli (all MDC) for valuable input on peptide clustering algorithms. This work was supported by the Deutsche Forschungsgemeinschaft (DFG, grant SE 1186/2-1), the National Natural Science Foundation of China (Grant No. 31861133013), and the Shenzhen Science and Technology Program (Grant No. KQTD20180411143432337)

## Author contributions

MS and WC initiated the project and acquired funding. AK, HZ, and MS designed the proteomic experiments. AK performed all proteomic experiments. HZ carried out the proteomic data analysis, developed the IsoFrac pipeline, and created the online protein isoform Shiny tool. QZ, MW and WC designed the transcriptomic experiments.QZ and MW conducted all transcriptomic experiments and data analysis, respectively. LF designed and performed the Western blot analysis of LDLR. MS and HZ wrote the manuscript with input from all authors.

## References

1. Bludau, I., and Aebersold, R. (2020). Proteomic and interactomic insights into the molecular basis of cell functional diversity. Nat. Rev. Mol. Cell Biol. 21, 327–340.

2. Buccitelli, C., and Selbach, M. (2020). mRNAs, proteins and the emerging principles of gene expression control. Nat. Rev. Genet. 21, 630–644.

3. Ule, J., and Blencowe, B.J. (2019). Alternative Splicing Regulatory Networks: Functions, Mechanisms, and Evolution. Mol. Cell 76, 329–345.

4. Mitschka, S., and Mayr, C. (2022). Context-specific regulation and function of mRNA alternative polyadenylation. Nat. Rev. Mol. Cell Biol. 10.1038/s41580-022-00507-5.

5. Blencowe, B.J. (2017). The Relationship between Alternative Splicing and Proteomic Complexity. Trends Biochem. Sci. 42, 407–408.

6. Smith, L.M., Agar, J.N., Chamot-Rooke, J., Danis, P.O., Ge, Y., Loo, J.A., Paša-Tolić, L., Tsybin, Y.O., Kelleher, N.L., and Consortium for Top-Down Proteomics (2021). The Human Proteoform Project: Defining the human proteome. Sci Adv 7, eabk0734.

7. Aebersold, R., Agar, J.N., Amster, I.J., Baker, M.S., Bertozzi, C.R., Boja, E.S., Costello, C.E., Cravatt, B.F., Fenselau, C., Garcia, B.A., et al. (2018). How many human proteoforms are there? Nat. Chem. Biol. 14, 206–214.

8. Hinnebusch, A.G., Ivanov, I.P., and Sonenberg, N. (2016). Translational control by 5’-untranslated regions of eukaryotic mRNAs. Science 352, 1413–1416.

9. Tress, M.L., Abascal, F., and Valencia, A. (2017). Alternative Splicing May Not Be the Key to Proteome Complexity. Preprint, 10.1016/j.tibs.2016.08.008 https://doi.org/10.1016/j.tibs.2016.08.008.

10. 10. Ezkurdia, I., Rodriguez, J.M., Carrillo-de Santa Pau, E., Vázquez, J., Valencia, A., and Tress, M.L. (2015). Most highly expressed protein-coding genes have a single dominant isoform. J. Proteome Res. 14, 1880–1887.

11. Toby, T.K., Fornelli, L., and Kelleher, N.L. (2016). Progress in Top-Down Proteomics and the Analysis of Proteoforms. Annu. Rev. Anal. Chem. 9, 499–519.

12. Kaulich, P.T., Cassidy, L., and Tholey, A. (2024). Identification of proteoforms by top-down proteomics using two-dimensional low/low pH reversed-phase liquid chromatography-mass spectrometry. Proteomics 24, e2200542.

13. Tran, J.C., Zamdborg, L., Ahlf, D.R., Lee, J.E., Catherman, A.D., Durbin, K.R., Tipton, J.D., Vellaichamy, A., Kellie, J.F., Li, M., et al. (2011). Mapping intact protein isoforms in discovery mode using top-down proteomics. Nature 480, 254–258.

14. Bekker-Jensen, D.B., Kelstrup, C.D., S Batth, T., Larsen, S.C., Haldrup, C., Bramsen, J.B., Sørensen, K.D., Høyer, S., Ørntoft, T.F., Andersen, C.L., et al. (2017). An Optimized Shotgun Strategy for the Rapid Generation of Comprehensive Human Proteomes. Cell Systems 4, 587–599.e4.

15. Nesvizhskii, A.I., and Aebersold, R. (2005). Interpretation of shotgun proteomic data: the protein inference problem. Mol Cell Proteomics 4, 1419–1440.

16. Umer, H.M., Audain, E., Zhu, Y., Pfeuffer, J., Sachsenberg, T., Lehtiö, J., Branca, R.M., and Perez-Riverol, Y. (2022). Generation of ENSEMBL-based proteogenomics databases boosts the identification of non-canonical peptides. Bioinformatics 38, 1470–1472.

17. Dou, Y., Yi, X., Olsen, L.K., and Zhang, B. (2022). SEPepQuant enables comprehensive protein isoform characterization in shotgun proteomics. bioRxiv, 2022.11.03.515027. 10.1101/2022.11.03.515027.

18. Bludau, I., Frank, M., Dörig, C., Cai, Y., Heusel, M., Rosenberger, G., Picotti, P., Collins, B.C., Röst, H., and Aebersold, R. (2021). Systematic detection of functional proteoform groups from bottom-up proteomic datasets. Nat. Commun. 12, 3810.

19. Dermit, M., Peters-Clarke, T.M., Shishkova, E., and Meyer, J.G. (2021). Peptide Correlation Analysis (PeCorA) Reveals Differential Proteoform Regulation. J. Proteome Res. 20, 1972–1980.

20. Miller, R.M., Jordan, B.T., Mehlferber, M.M., Jeffery, E.D., Chatzipantsiou, C., Kaur, S., Millikin, R.J., Dai, Y., Tiberi, S., Castaldi, P.J., et al. (2022). Enhanced protein isoform characterization through long-read proteogenomics. Genome Biol. 23, 69.

21. Sinitcyn, P., Richards, A.L., Weatheritt, R.J., Brademan, D.R., Marx, H., Shishkova, E., Meyer, J.G., Hebert, A.S., Westphall, M.S., Blencowe, B.J., et al. (2023). Global detection of human variants and isoforms by deep proteome sequencing. Nat Biotechnol 41, 1776–1786.

22. Kurzawa, N., Leo, I.R., Stahl, M., Kunold, E., Becher, I., Audrey, A., Mermelekas, G., Huber, W., Mateus, A., Savitski, M.M., et al. (2023). Deep thermal profiling for detection of functional proteoform groups. Nat Chem Biol 19, 962–971.

23. Chen, C., Wen, M., and Jin, Y. (2022). 1DE-MS Profiling for Proteoform-Correlated Proteomic Analysis, by Combining SDS-PAGE, Whole-Gel Slicing, Quantitative LC–MS/MS, and Reconstruction of Gel Distributions of Several Thousands of Proteins. J. Proteome Res. 10.1021/acs.jproteome.2c00180.

24. Thiede, B., Treumann, A., Kretschmer, A., Söhlke, J., and Rudel, T. (2005). Shotgun proteome analysis of protein cleavage in apoptotic cells. Proteomics 5, 2123–2130.

25. Wang, X., You, X., Langer, J.D., Hou, J., Rupprecht, F., Vlatkovic, I., Quedenau, C., Tushev, G., Epstein, I., Schaefke, B., et al. (2019). Full-length transcriptome reconstruction reveals a large diversity of RNA and protein isoforms in rat hippocampus. Nature Communications 10, 1–15.

26. Cox, J., and Mann, M. (2008). MaxQuant enables high peptide identification rates, individualized p.p.b.-range mass accuracies and proteome-wide protein quantification. Nat. Biotechnol. 26, 1367–1372.

27. Perez-Riverol, Y., Xu, Q.-W., Wang, R., Uszkoreit, J., Griss, J., Sanchez, A., Reisinger, F., Csordas, A., Ternent, T., Del-Toro, N., et al. (2016). PRIDE Inspector Toolsuite: Moving Toward a Universal Visualization Tool for Proteomics Data Standard Formats and Quality Assessment of ProteomeXchange Datasets. Mol Cell Proteomics 15, 305–317.

28. Perez-Riverol, Y., Bandla, C., Kundu, D.J., Kamatchinathan, S., Bai, J., Hewapathirana, S., John, N.S., Prakash, A., Walzer, M., Wang, S., et al. (2025). The PRIDE database at 20 years: 2025 update. Nucleic Acids Res 53, D543–D553.

29. Mann, M. (2020). The Origins of Organellar Mapping by Protein Correlation Profiling. Proteomics 20, e1900330.

30. Kristensen, A.R., Gsponer, J., and Foster, L.J. (2012). A high-throughput approach for measuring temporal changes in the interactome. Preprint, 10.1038/nmeth.2131 https://doi.org/10.1038/nmeth.2131.

31. Imami, K., Milek, M., Bogdanow, B., Yasuda, T., Kastelic, N., Zauber, H., Ishihama, Y., Landthaler, M., and Selbach, M. (2018). Phosphorylation of the Ribosomal Protein RPL12/uL11 Affects Translation during Mitosis. Mol. Cell 72, 84–98.e9.

32. Dengjel, J., Høyer-Hansen, M., Nielsen, M.O., Eisenberg, T., Harder, L.M., Schandorff, S., Farkas, T., Kirkegaard, T., Becker, A.C., Schroeder, S., et al. (2012). Identification of Autophagosome-associated Proteins and Regulators by Quantitative Proteomic Analysis and Genetic Screens. Preprint, 10.1074/mcp.m111.014035 https://doi.org/10.1074/mcp.m111.014035.

33. Christoforou, A., Mulvey, C.M., Breckels, L.M., Geladaki, A., Hurrell, T., Hayward, P.C., Naake, T., Gatto, L., Viner, R., Martinez Arias, A., et al. (2016). A draft map of the mouse pluripotent stem cell spatial proteome. Nat. Commun. 7, 8992.

34. Aryal, U.K., Xiong, Y., McBride, Z., Kihara, D., Xie, J., Hall, M.C., and Szymanski, D.B. (2014). A proteomic strategy for global analysis of plant protein complexes. Plant Cell 26, 3867–3882.

35. Dunkley, T.P.J., Watson, R., Griffin, J.L., Dupree, P., and Lilley, K.S. (2004). Localization of Organelle Proteins by Isotope Tagging (LOPIT). Preprint, 10.1074/mcp.t400009-mcp200 https://doi.org/10.1074/mcp.t400009-mcp200.

36. Andersen, J.S., Wilkinson, C.J., Mayor, T., Mortensen, P., Nigg, E.A., and Mann, M. (2003). Proteomic characterization of the human centrosome by protein correlation profiling. Nature 426, 570–574.

37. 37. Foster, L.J., de Hoog, C.L., Zhang, Y., Zhang, Y., Xie, X., Mootha, V.K., and Mann, M. (2006). A mammalian organelle map by protein correlation profiling. Cell 125, 187–199.

38. Laemmli, U.K. (1970). Cleavage of structural proteins during the assembly of the head of bacteriophage T4. Nature 227, 680–685.

39. Witkowski, C., and Harkins, J. (2009). Using the GELFREE 8100 Fractionation System for molecular weight-based fractionation with liquid phase recovery. J. Vis. Exp. 10.3791/1842.

40. Müller, T., Kalxdorf, M., Longuespée, R., Kazdal, D.N., Stenzinger, A., and Krijgsveld, J. (2020). Automated sample preparation with SP3 for low-input clinical proteomics. Mol. Syst. Biol. 16, e9111.

41. Zecha, J., Satpathy, S., Kanashova, T., Avanessian, S.C., Kane, M.H., Clauser, K.R., Mertins, P., Carr, S.A., and Kuster, B. (2019). TMT Labeling for the Masses: A Robust and Cost-efficient, In-solution Labeling Approach. Mol. Cell. Proteomics 18, 1468–1478.

42. Grzybowska, E.A. (2012). Human intronless genes: functional groups, associated diseases, evolution, and mRNA processing in absence of splicing. Biochem. Biophys. Res. Commun. 424, 1–6.

43. Andreev, D.E., Loughran, G., Fedorova, A.D., Mikhaylova, M.S., Shatsky, I.N., and Baranov, P.V. (2022). Non-AUG translation initiation in mammals. Genome Biol 23, 111.

44. Manjunath, L.E., Singh, A., Som, S., and Eswarappa, S.M. (2023). Mammalian proteome expansion by stop codon readthrough. Wiley Interdiscip Rev RNA 14, e1739.

45. Rogers, L.D., and Overall, C.M. (2013). Proteolytic post-translational modification of proteins: proteomic tools and methodology. Mol Cell Proteomics 12, 3532–3542.

46. Campello, R.J.G.B., Campello, R.J.G., Moulavi, D., and Sander, J. (2013). Density-Based Clustering Based on Hierarchical Density Estimates. Preprint, 10.1007/978-3-642-37456-2_14 https://doi.org/10.1007/978-3-642-37456-2_14.

47. Currey, L., Thor, S., and Piper, M. (2021). TEAD family transcription factors in development and disease. Development 148. 10.1242/dev.196675.

48. Melby, J.A., Brown, K.A., Gregorich, Z.R., Roberts, D.S., Chapman, E.A., Ehlers, L.E., Gao, Z., Larson, E.J., Jin, Y., Lopez, J.R., et al. (2023). High sensitivity top-down proteomics captures single muscle cell heterogeneity in large proteoforms. Proc Natl Acad Sci U S A 120, e2222081120.

49. Shi, Y., Mowery, R.A., Ashley, J., Hentz, M., Ramirez, A.J., Bilgicer, B., Slunt-Brown, H., Borchelt, D.R., and Shaw, B.F. (2012). Abnormal SDS-PAGE migration of cytosolic proteins can identify domains and mechanisms that control surfactant binding. Protein Sci. 21, 1197–1209.

50. Burgess, R.R. (2009). Important but little known (or forgotten) artifacts in protein biochemistry. Methods Enzymol. 463, 813–820.

