## Supplemental Figures for "An integrated landscape of mRNA and protein isoforms"

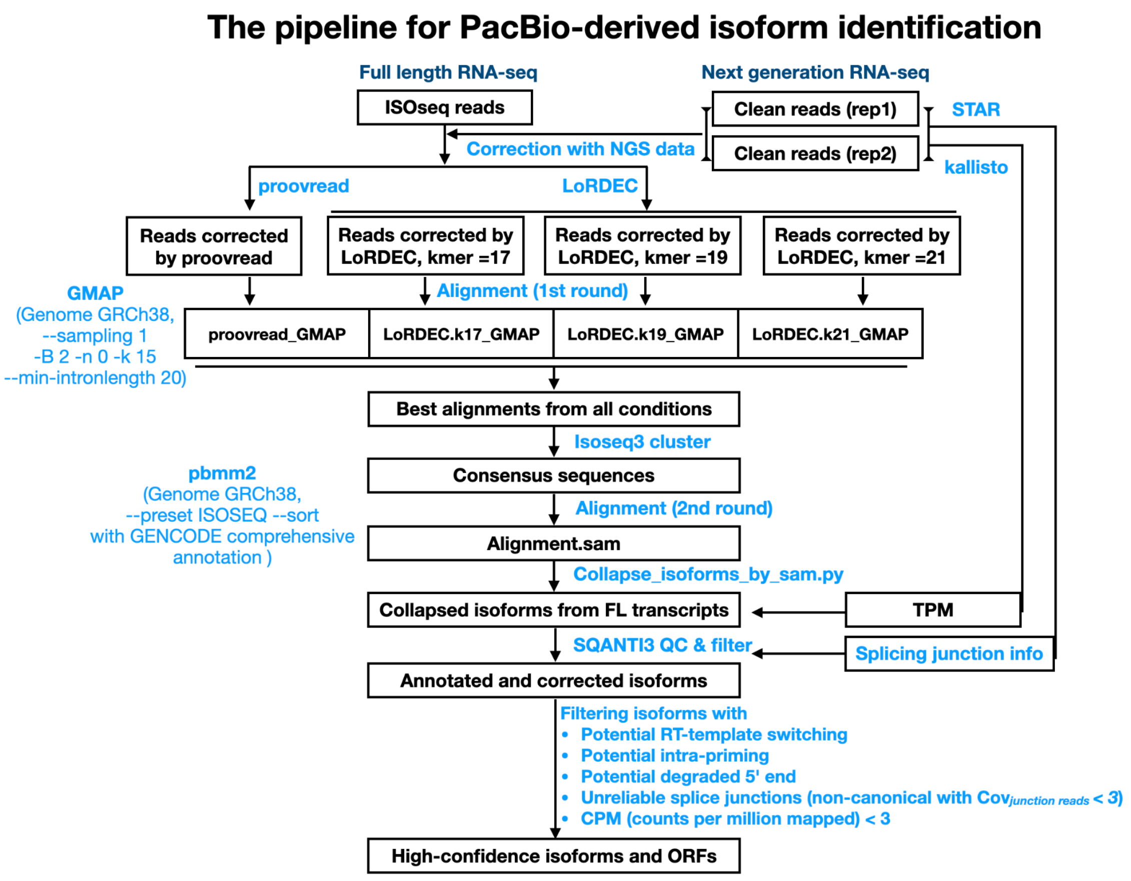


**Supplemental Figure 1**
The bioinformatics pipeline for identifying full-length transcripts using PacBio ISO-seq and NGS RNA-seq data.

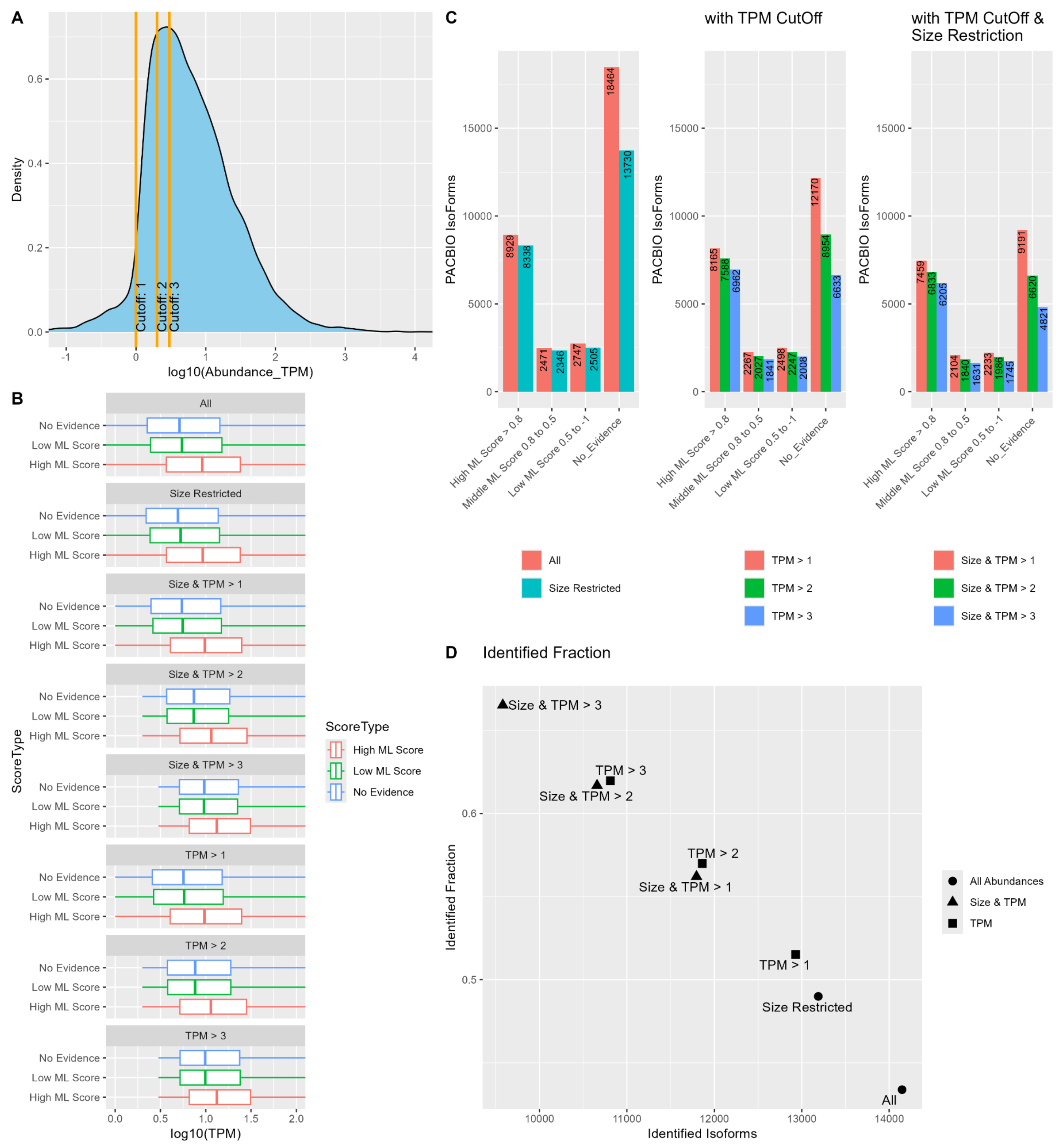


**Supplemental Figure 2**

Exclusive Assignment of PacBio isoforms to IsoFrac protein isoforms. A) TPM Abundance distribution of RNA Transcripts. Different tested cutoffs of abundance are colored in orange. B) Transcript abundance boxplots for different Cutoff subsets. Data is further binned into three groups based on Isofrac identification confidence score “high” (score > 0.8), “middle” (score between 0.8 and 0.5), “low” (< 0.5) and “No evidence”). C) Counts of PacBIO isoforms for the different Isofrac cross mapping bins without cutoffs (left), with TPM cutoff (middle) and with TPM and size restriction cutoff (right). The “no evidence” fraction is overproportionally affected by these cutoffs. D) Counts and the corresponding fraction of protein isoforms for different cutoffs identified in both, proteomics and transcriptomics.

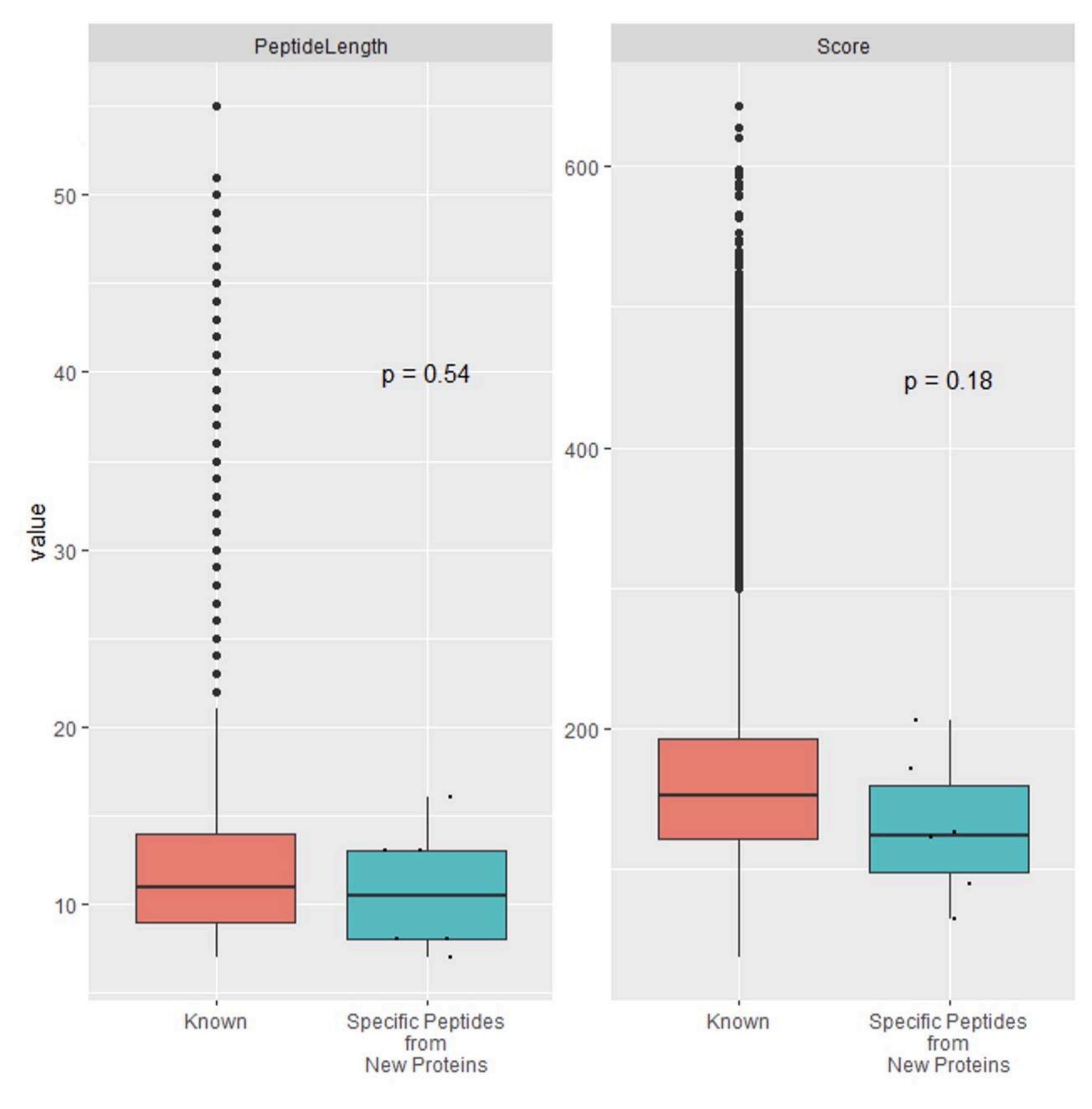


**Supplemental Figure 3**
Length and Score distributions of known and newly identified peptides. The p-value of a Wilcoxon test statistic is depicted in the figure.

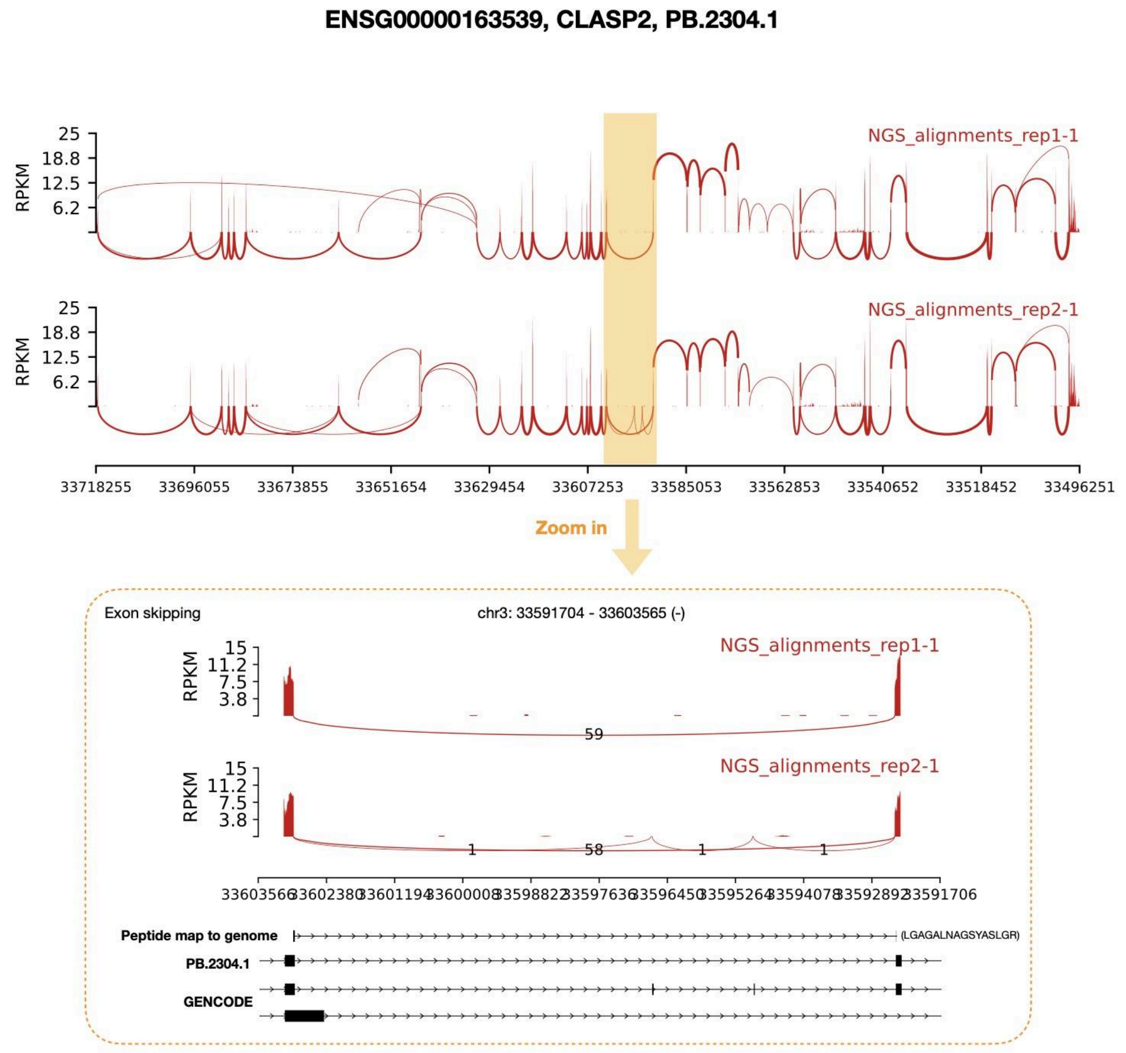


**Supplemental 4.1**

Sashimi Plot shows that NGS sequencing, PacBio sequencing, and peptide segments effectively support the newly identified transcript of the gene CLASP2 (ENSG00000163539). Upper: Sashimi Plot displays the coverage of short-read NGS sequencing and the pattern of splice junctions. Bottom: A zoom-in on the region which peptide alignment to the transcriptome. The numbers on the arcs of the Sashimi Plot represent the read counts corresponding to the respective splice junctions, while the bottom of the Sashimi Plot shows the alignment of the peptide segments, as well as the newly discovered transcript by PacBio and the known transcripts from GENCODE.

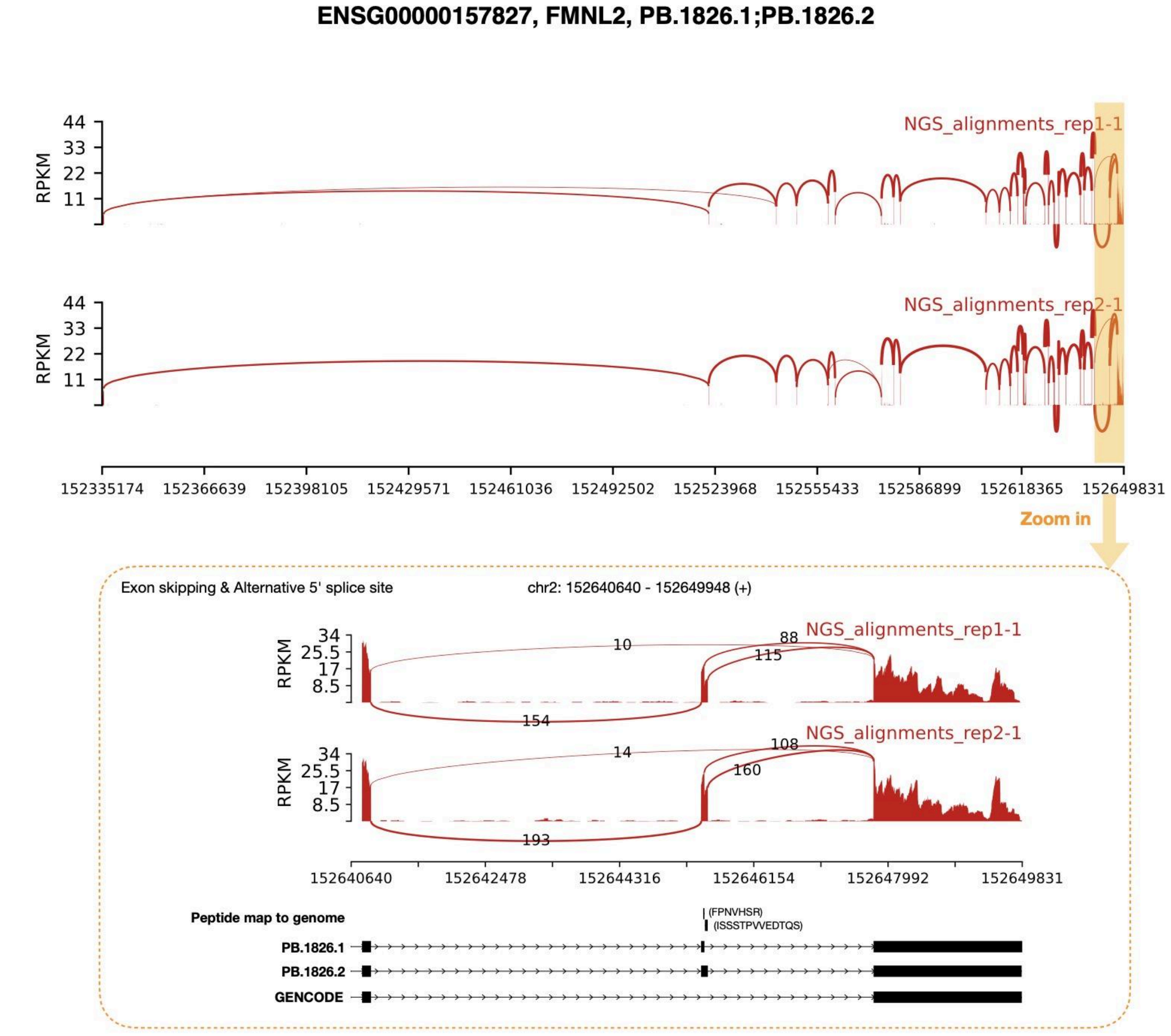


**Supplemental 4.2**
Sashimi Plot for FMNL2 (ENSG00000157827) effectively supporting the newly identified transcript. For details see Supplemental Figure 4.1.

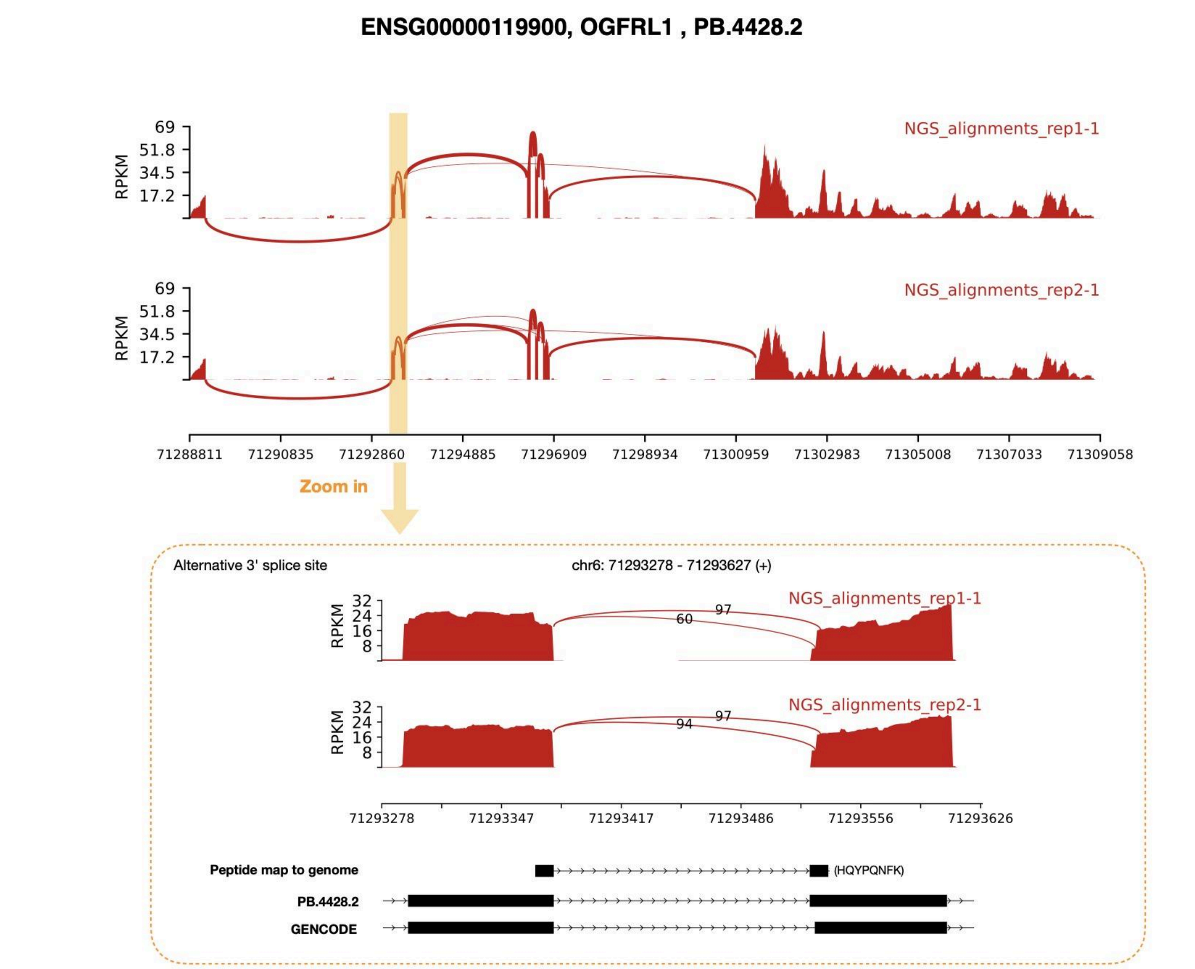


**Supplemental 4.3**
Sashimi Plot for OGFRL1 (ENSG00000119900) effectively supporting the newly identified transcript. For details see Supplemental Figure 4.1.

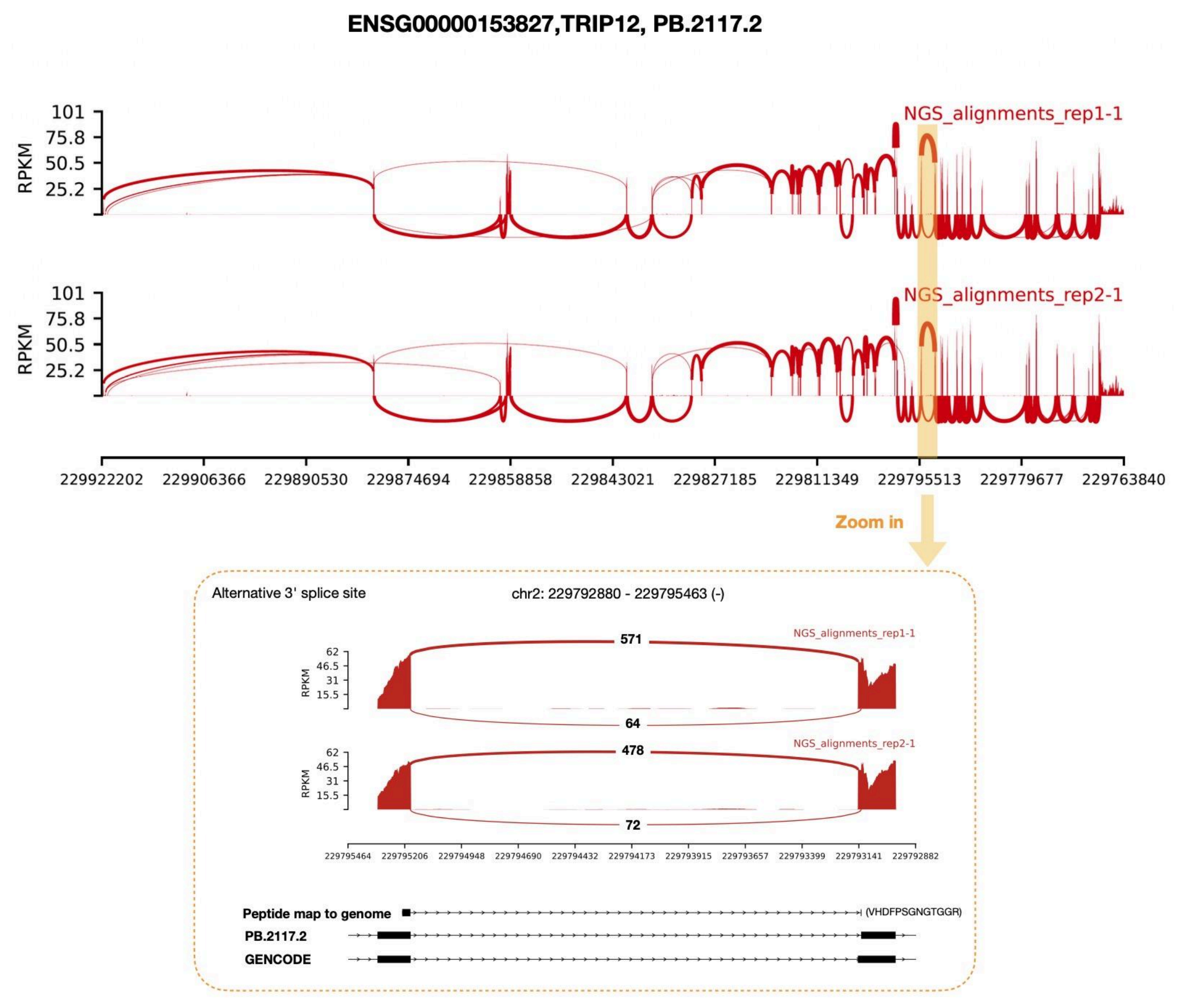


**Supplemental 4.4**
Sashimi Plot for TRIP12 (ENSG00000153827) effectively supporting the newly identified transcript. For details see Supplemental Figure 4.1.

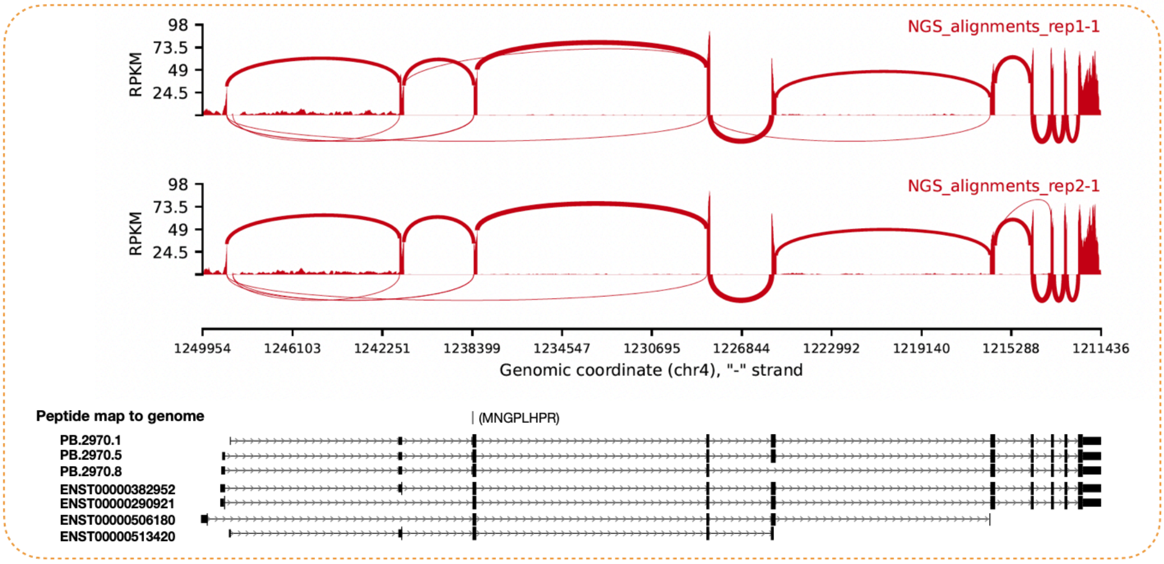


**Supplemental 4.5**
Sashimi Plot for CTBP1 (ENSG00000159692) effectively supporting the identified alternative translation start site for three PacBio transcripts. For details see Supplemental Figure 4.1.

**CTTNBP2NL**

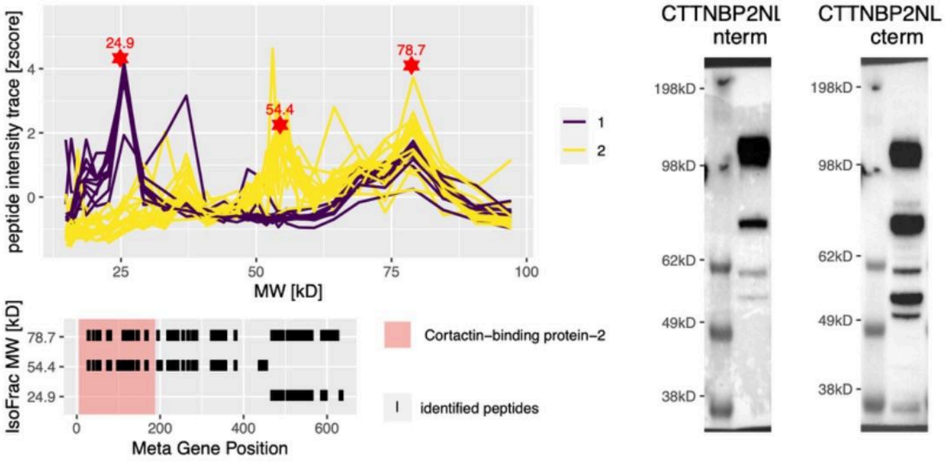


**NELFA**

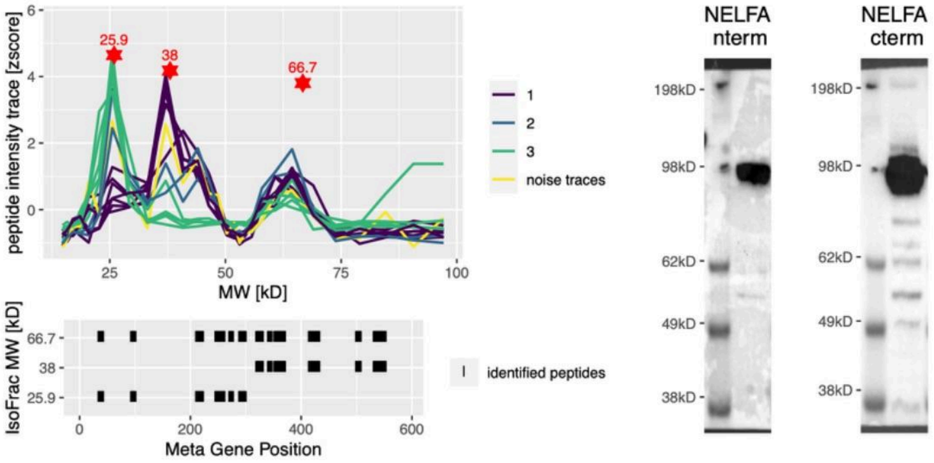


**RRP1B**

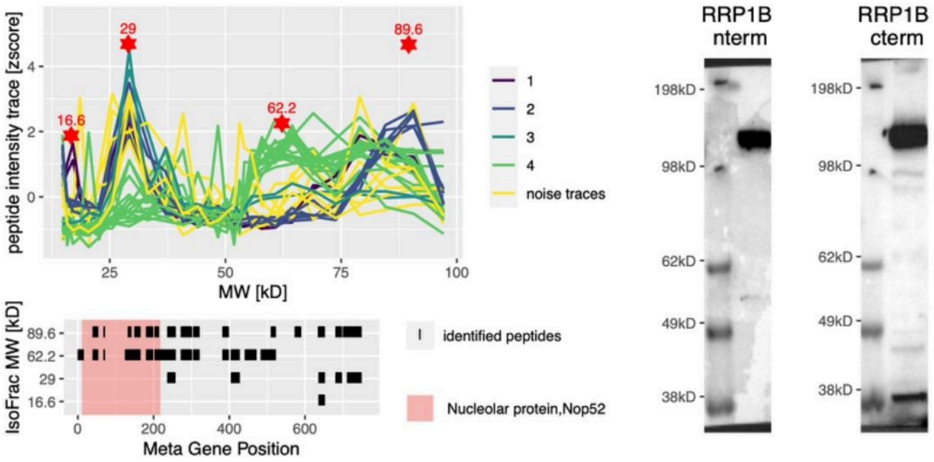

**Supplemental Figure 5**
Validation of proteomic cleavage derived isoforms for CTTNBP2NL, NELFA and RRP1B. For each gene, the peptide traces across fractions with their corresponding identified isoform candidates (red stars, upper left subpanel) are shown. After manual validation of each automatically assigned peptide set, these peptides were mapped to the canonical protein sequence of each gene (lower left subpanel). Peptides are indicated as black segments and known protein domains are marked by background color. Western blots of n-terminal or c-terminal tagged constructs of each genes are shown in the right hand subpanels. Please note that the mass added by the tag to the total protein mass is ~30kD.
